# CD44v6 expression is a novel predictive marker of therapy response and poor prognosis in gastric cancer patients

**DOI:** 10.1101/468934

**Authors:** Carla Pereira, Daniel Ferreira, Carolina Lemos, Diana Martins, Nuno Mendes, Daniela Almeida, Pedro Granja, Fátima Carneiro, Raquel Almeida, Gabriela M Almeida, Carla Oliveira

**Affiliations:** i3S - Instituto de Investigação e Inovação em Saúde, Universidade do Porto, Porto, Portugal; Ipatimup - Institute of Molecular Pathology and Immunology of the University of Porto, Porto, Portugal; INEB, Instituto de Engenharia Biomédica, Universidade do Porto, Porto, Portugal; IBMC - Institute for Molecular and Cell Biology, Porto, Portugal; ICBAS - Instituto Ciências Biomédicas Abel Salazar, Universidade do Porto, Porto, Portugal; Medical Oncology Department, Centro Hospitalar de São João, Porto, Portugal; Faculty of Medicine of the University of Porto, Porto, Portugal; Department of Pathology, Centro Hospitalar de São João, Porto, Portugal; Faculty of Sciences of the University of Porto, Porto, Portugal

**Author notes:** Correspondence to: C. Oliveira, Expression Regulation in Cancer Group, Ipatimup, Instituto de Investigação e Inovação em Saúde (I3S), Rua Alfredo Allen, N° 208, 4200-135 Porto Portugal; or G.M. Almeida, Expression Regulation in Cancer Group, Ipatimup, Instituto de Investigação e Inovação em Saúde (I3S), Rua Alfredo Allen, N° 208, 4200-135 Porto Portugal. Both CP and DF contributed equally to the work.

**Keywords:** gastric cancer, CD44v6, prognostic marker, predictive marker, therapy response, gastric cancer survival, tumor heterogeneity

## Abstract

Late diagnosis, modest treatment options and lack of predictive markers of therapy response dictate the poor overall survival (OS) of ∼1 year in most gastric cancer (GC) patients. We hypothesized that the level of CD44v6 expression in tumor cells could predict therapy response and prognosis in GC patients.

We analyzed a surgical tumor series of GC patients for the extension of CD44v6 membranous immuno-expression, clinical-pathological features, patient survival, and response to therapy. By integrating this information, we assessed the value of CD44v6 expression to predict benefit from current treatment regimens and prognosis in GC patients. We used GC cell lines and mouse xenografts to assess and/validate the biological impact of CD44v6 expression in GC cells behavior.

We demonstrated that GC patients whose tumors present higher levels of CD44v6 membranous expression benefit from adding chemotherapy to surgery as opposed to those without CD44v6 expression. Moreover, patients bearing CD44_high tumors presented worse OS than those bearing CD44_absent/low tumors, consolidating the role of CD44v6 expression as an independent factor of poor prognosis in this disease. Finally, our *in vitro* and patients’ data pinpoints the CD44v6+ cell population as the driver of tumor recurrence following conventional chemotherapy, in heterogeneous tumors composed by CD44v6- and CD44v6+ cells.

Our study pioneers the identification of CD44v6 as a potential predictive marker of response to conventional chemotherapy, and consolidates CD44v6 as an independent marker of poor prognosis in GC. Overall, our data strongly supports selection of patients with high CD44v6 expressing tumors for conventional chemotherapy with or without surgery, regardless of the TNM stage.

## Introduction

Gastric cancer (GC) is the 3^rd^ leading cause of cancer related mortality worldwide, with >750,000 deaths estimated to occur every year.^1^Over 70% of GC patients present with locally advanced and/or unresectable disease, for whom conventional chemotherapy (mostly platinum-based) becomes the main treatment option, with a median overall survival (OS) of ∼1 year.^2,3^Despite treatment improvements, the use of targeted therapies in GC has proved disappointing,^4^with the only approved therapies (Trastuzumab, against HER2 and Ramucirumab, against VEGFR2) showing limited OS improvement (2 to 3 months).^5,6^Therefore, it is crucial to identify biomarkers that can better define prognosis, but mainly that allow predicting who is likely benefitting more from a given treatment regimen. This would trigger better patient stratification for treatment, likely improving OS.

CD44, the main hyaluronic acid receptor, is a transmembrane glycoprotein involved in key cellular processes, such as lymphocyte activation, recirculation and homing, as well as epithelial cell adhesion and migration.^7,8^The human CD44 gene (NG_008937) encodes a polymorphic group of proteins generated by alternative splicing. The standard CD44 isoform (CD44s) includes only the constitutive exons, while the variant isoforms (CD44v) contain one or more variable exons (in addition to the constitutive ones).^7-9^CD44s is ubiquitously expressed at the surface of most mammalian cells, whereas the expression of CD44v is highly restricted (e.g. during lymphocyte activation and homing) and to specific tissue types.^7-10^Aberrant expression of CD44v isoforms may occur in diseased cells, and have been associated particularly with several cancer-associated features like invasion and metastization.^9,11,12^Moreover, CD44 and its CD44v isoforms are expressed as surface markers of cancer stem cells (CSCs), influencing key CSC-associated properties such as tumor initiation, self-renewal, metastasis and chemoresistance.^9,13-15^

In the stomach, we have shown that CD44s isoform is widely expressed in both normal and diseased gastric epithelial cells.^16^In contrast, we showed that CD44v6-containing isoforms, which are absent from normal gastric epithelial cells, become overexpressed in stomach premalignant and malignant lesions and display high expression in ∼70% of all GCs.^16^

Meta-analysis data showed that positive CD44 expression was significantly associated with worse GC patients’ prognosis and lower OS, and suggested a similar association for CD44v6 expression.^17^Comparable data exists for other cancer types such as lung, colorectal and breast.^13^Recently, two reports showed a positive association between increased expression of CD44v6 and worse OS in GC patients.^18,19^However, both studies showed a positive correlation between CD44v6 expression and higher TNM staging, leaving reasonable doubt as to whether CD44v6 tumor expression is an independent factor of poor prognosis in GC. In addition, most articles published on CD44v6 in GC cohorts included a limited number of patients, making it difficult to obtain statistically significant results. No reports exist, exploring whether tumor CD44v6 expression influences patient therapy response.

Therefore, we aimed to clarify the role of CD44v6 in GC by using a large GC cohort to investigate, simultaneously, the relationship between CD44v6 expression and clinical-pathological features, patient OS and therapy response.

## Materials and Methods

All reagents were purchased from ThermoFisher Scientific (Waltham, MA, USA) unless otherwise stated.

### Patient sample/data collection and tissue microarray (TMA) preparation

A cohort of 334 GC patients surgically treated at Centro Hospitalar de São João (CHSJ), Portugal, and part of the Tumor biobank of CHSJ/Ipatimup^20^was assembled (n=334) (Appendix Fig 1). A TMA was prepared from paraffin-embedded tumor material and patients’ clinical-pathological, treatment and follow-up data collected. This study was approved by the Institutional Ethics Committee of CHSJ (Ethics Committee references CES 122/15 and CES 117/18), and informed patient consent obtained. This study is REMARK compliant.

### CD44v6 immunohistochemistry of GC samples

Immunohistochemistry (IHC) staining for CD44v6 (clone MA54) was performed in 3-µm sections from TMAs, using the automated Ventana BenchMark ULTRAStaining System, using the OptiView DAB IHC Detection Kit (both from Roche/Ventana Medical Systems, Tucson, AZ, USA) according to manufacturers’ instructions. Positive (human skin) and negative staining controls were performed in parallel with paraffin sections. The percentage of tumor cells displaying membranous expression of CD44v6 was assessed, for each sample, by four researchers in two independent evaluations performed in a blind manner. Four categories were defined to classify the extent of CD44v6 tumor expression: “CD44_0” – no staining at the cell membrane; “CD44_1+” – membranous staining in less than 10% of tumor cells; “CD44_2+” – membranous staining in between 10 and 50% of tumor cells; “CD44_3+” – membranous staining in more than 50% of the tumor cells (Fig 1A).

**Fig 1:**
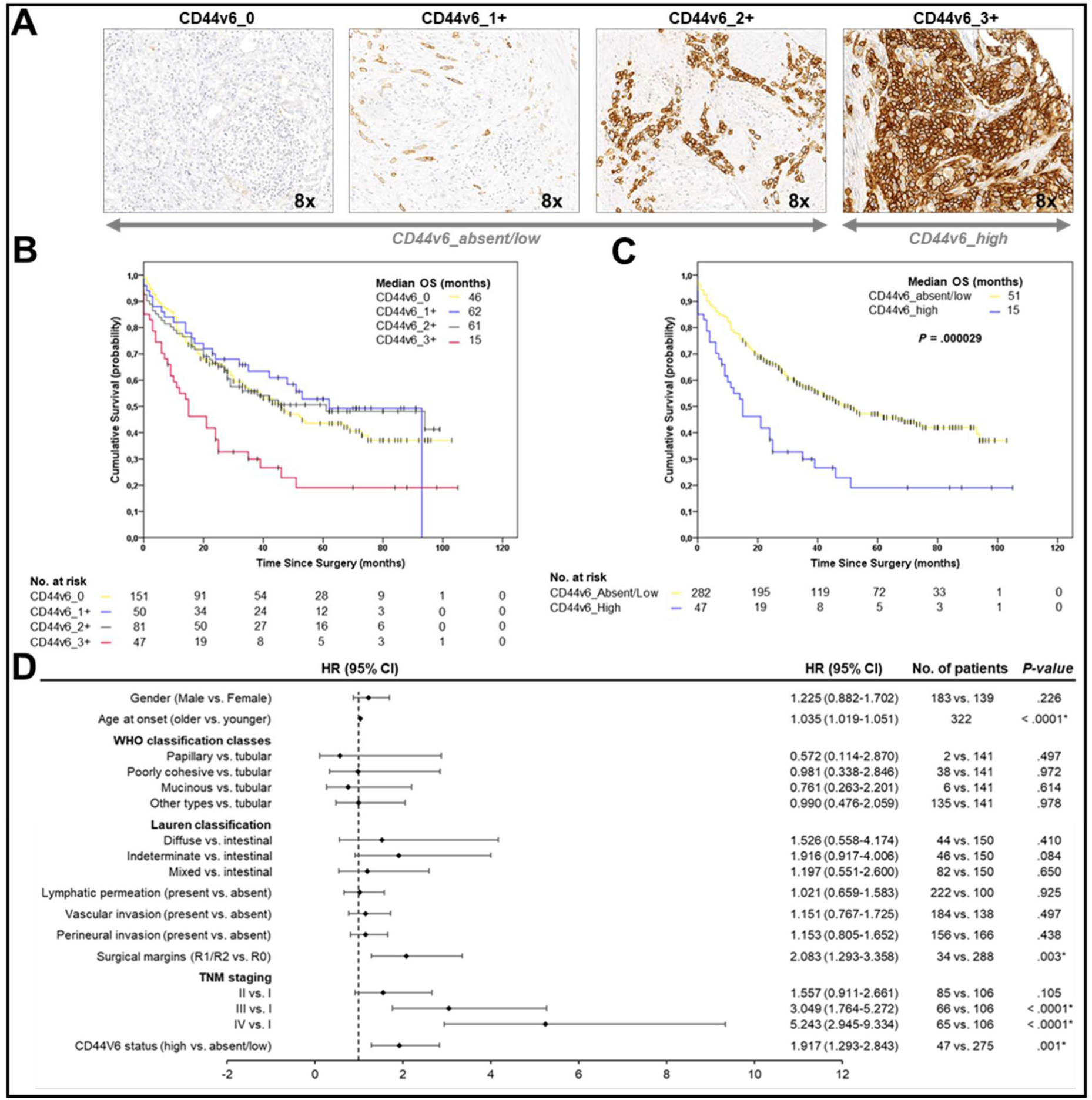
(A) Immunohistochemistry characterization of samples regarding the presence and extent of CD44v6 expression. Images are representative of the four categories defined: CD44_0 (no staining at the cell membrane); CD44_1+ (membranous staining in < 10% of tumor cells); CD44_2+ (membranous staining in ≥ 10% and < 50% of tumor cells); CD44_3+ (membranous staining in ≥ 50% of tumor cells); **(B)** Kaplan-Meier estimates showing OS of GC patients according to CD44v6 sub-categories and corresponding median OS - CD44_3+ patients have worse OS compared to other sub-categories (*P* < .001); **(C)** Kaplan-Meier estimates showing OS of GC patients according to absent/low and high expression of CD44v6 and corresponding median OS – “CD44_High” patients have the worst prognosis (*P* < .0001); **(D)** Forest plot of the multivariate analysis. Expression of CD44v6 in more than 50% of tumor cells was identified as an independent factor of poor prognosis in GC.

### Cell lines and culture conditions

Human GC cell lines MKN74 and MKN45 were purchased from the JCRB Cell Bank (Japanese Collection of Research Bioresources Cell Bank), and the non-commercial cell line GP202 cell line was established at Ipatimup.^21^All cell lines were cultured in RPMI 1640 medium with 10% heat inactivated fetal bovine serum (FBS) (Biowest, Nuaillé, France). Cell lines were maintained at 37 °C and 5% CO_2_ in a high humidity atmosphere and sub-cultured every 3 to 4 days. Cells were grown in the absence of antibiotics except for cell selection in MKN74 cells, where G418 was used. Cells were never continuously cultured for more than 4 months. Cell identification was confirmed by STR analysis and cells were confirmed to be free of mycoplasma contamination.

### Generation of an isogenic cell line model of tumor CD44v6 status

Sequences for CD44s and CD44v6 (a v3-v10 transcript present in GP202) were cloned into a pIRES-EGFP2 plasmid.^22^MKN74 cells were transfected with either the empty vector (MKN74_Mock), pIRES-EGFP2_CD44v6 (MKN74_CD44v6) or pIRES-EGFP2_CD44s (MKN74_CD44s) and pure cell populations obtained by selective pressure with 1 mg/mL of G418 and bead sorting with the Magnetic Separation kit CELLection Pan Mouse IgG kit from Dynabeads^®^, performed according to the manufacturers’ instructions.

### Assessment of cell survival upon drug treatments

MKN74_Mock, MKN74_CD44v6 and MKN74_CD44s cells were seeded in 96-well plates (2.5×10^3^per well) under normal conditions (5% CO_2_humidified atmosphere at 37 °C) and allowed to adhere for 24 h. Cells were then incubated with different concentrations of cisplatin or 5-FU for 48 h and processed for the Sulforhodamine B (SRB) assay. Briefly, at the required times, cells were fixed in 10% trichloroacetic acid (TCA) for 1 h on ice, proteins stained with 4% SRB solution (Sigma-Aldrich; Poole, UK) for 30 min, wells washed repeatedly with 1% acetic acid to remove the unbound dye, and the protein-bound stain was solubilized with 10 mM Tris solution. The SRB absorbance was measured at 560 nm and background corrected at 655 nm, using a microplate reader (PowerWave HT Microplate Spectrophotometer; BioTek, Bad Friedrichshall, Germany). At least three independent experiments were performed, each measured in triplicate. Cell survival for each drug treatment was calculated, as a percentage, in relation to the respective vehicle-treated control, for each cell line.

### Assessment of cisplatin-induced apoptosis

MKN74_Mock, MKN74_CD44v6 and MKN74_CD44s cells were seeded in 6-well plates (1.0×10^5^cells/ well), left to adhere for 24 h, and incubated with 10 µM of cisplatin for 24 and 48 h. Cisplatin-induced apoptosis was assayed by labelling cells with Annexin V-APC antibody (ImmunoTools, Friesoythe, Germany), according to the manufacturer’s instructions. Measurement of phosphatidylserine externalization was analyzed using an Accuri C6 flow cytometer and Accuri C6 software (BD Biosciences, Franklin Lakes, NJ, USA), plotting at least 20,000 events per sample. Results represent the average of at least three independent experiments.

### CD44v6 expression inhibition by siRNA

GP202 and MKN45 cells were transfected with siRNAs for CD44v6 using Lipofectamine^®^ RNAiMax Transfection Reagent, according to the manufacturers’ instructions. Briefly, cells were seeded in 6 well-plates (2×10^5^cells/well and 1.5×10^5^cells/well, respectively GP202 and MKN45 cell lines). After 24 h, lipid based conjugates were prepared by mixing 10 nM of non-targeting siRNA (negative control DS NC1; iDT, Leuven, Belgium), or human CD44v6 siRNA (Sense strand: 5’-GCGUCAGGUUCCAUAGGAAUCCUTT −3’ and Antisense strand: 5’-AAAGGAUUCCUAUGGAACCUGACGCAG −3’, custom made from iDT, Leuven, Belgium), to diluted Lipofectamine^®^ RNAiMax Reagent in 1:1 ratio. The conjugates were incubated for 5 min at room temperature and added dropwise to the cells. Upon 48 h of incubation, protein extraction and Western blots were performed to evaluate the efficacy of CD44v6 silencing. In addition, *in vitro* cisplatin treatments were carried out and apoptosis assessed (as described above).

### Percentage of CD44v6+ cells upon cisplatin incubation

MKN74_Mock and MKN74_CD44v6 were mixed and seeded in 6 well plates with an initial density of 5×10^4^cells/well (50:50 proportion), allowed to grow for 24 h and incubated with 10 μM of cisplatin or vehicle (0.9% (v/v) NaCl) for 6 h. Cells were then washed with PBS, growth medium was replaced with cisplatin-free media and co-cultures were maintained during 15 days. CD44v6 cell enrichment was assessed at several time points by flow cytometry using a CD44v6 conjugated antibody. Briefly, at each time point, cells were washed with PBS, versinized for 15 min, blocked in PBS - 2% FBS for 30 min and incubated with CD44v6-APC conjugated antibody (2F10; 10 μL/106 cells; 30 min; R&D Systems, McKinley Place, MN, USA). Cells were subsequently washed with PBS - 2% FBS and immediately proceeded for analysis. Fluorescence was analyzed using a FACS Canto II (BD Biosciences, Franklin Lakes, NJ, USA). The mean fluorescence intensity was measured for at least 20,000 gated events per sample and data analyzed using the software FlowJo, version 10.

### Statistical Analysis

Age, gender and patients’ clinical-pathological features were compared according to CD44v6 expression and treatment type. Univariate analyses for categorical variables were performed using Chi-square or Fisher-exact test as appropriate. Continuous variables were analyzed using Student’s t-test or One-way ANOVA. For the statistical analyses of experiments in Fig 2 a Two-way ANOVA was performed. Post-hoc tests were performed for significant One- or Two-way ANOVA results using Tukey’s Post Hoc Test.

**Fig 2:**
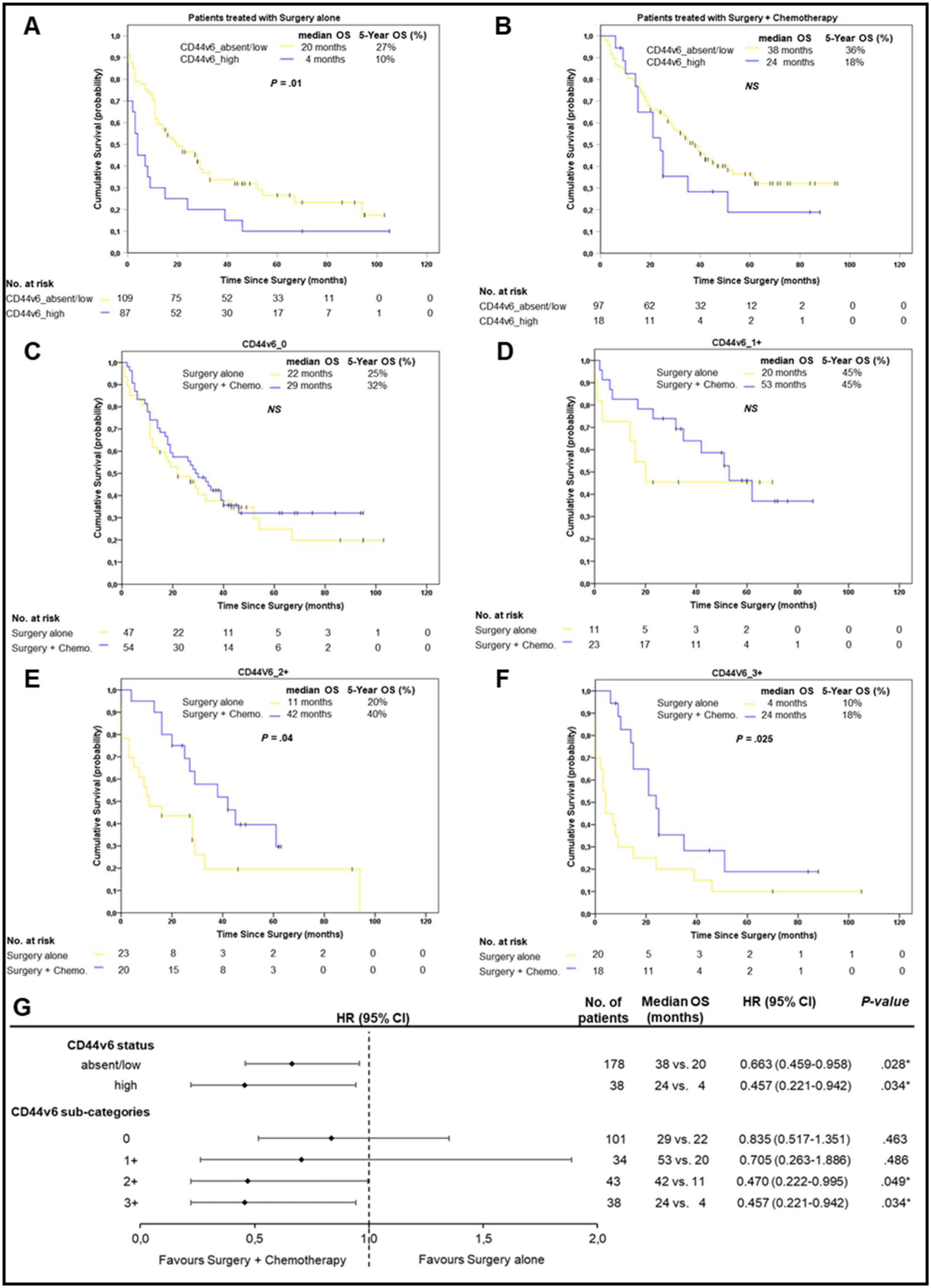
(A) Kaplan-Meier estimates showing OS of patients treated with surgery alone or **(B)** surgery plus chemotherapy, according to CD44v6 status; **(C)** Kaplan-Meier estimates showing OS of CD44_0 **(D)** CD44_1+, **(E)** CD44_2+ and **(F)** CD44_3+ GC patients, according to whether they were treated with surgery alone or with surgery plus chemotherapy; **(G)** Forest plot showing the benefit of treating patients with surgery plus chemotherapy according to CD44v6 classification.

Kaplan-Meier estimates of Overall Survival (OS) and Relapse-Free Survival (RFS) were obtained between the groups. Afterwards, Multivariate Cox regression analysis of OS and RFS were performed, and the hazard ratio (HR) and 95% CI estimated, in order to determine factors that were independently associated with RFS and OS. A *P*-value < .05 was considered significant and all analyses were performed using IBM SPSS Statistics version 24 for Windows. This study is TRIPOD compliant.

## Results

### Consolidation of CD44v6 *de novo* expression in tumor cells as a poor prognosis marker in GC patients

Data on clinical-pathological features and treatment type (Appendix Table 1), OS and RFS was collected from a surgical series of 334 GC patients. This series has a good representation of all disease stages and, as expected, patient OS and RFS significantly worsens with increasing pTNM (Pathological Tumor-Node-Metastasis) staging (Appendix Fig 2A and B). Analysis of OS and RFS in chemotherapy treated *vs*. untreated stage II to IV patients, demonstrated that chemotherapy plus surgery increases OS (*P* < .005), although no advantage was observed for RFS (Appendix Fig 3A and B).

The extent of CD44v6 membranous expression was analyzed in all cases and tumors were classified into four sub-categories, from absent to high (0, 1+, 2+ and 3+; Fig 1A): 46% (155/334) were CD44_0; 15% (50/334) were CD44_1+; 25% (82/334) were CD44_2+, and; 14% (47/334) were CD44_3+. We found a strong association between high CD44v6 expression and poorer OS (Fig 1B). Indeed, GC patients whose tumors express CD44v6 in >50% of tumor cells (CD44_3+), have statistically significant worse OS than those lacking CD44v6 expression or if CD44v6 is present in <50% of tumor cells (*P* < .001) (Fig 1B). Therefore, it seemed reasonable to group patients into two expression categories: “CD44_absent/low” (combining 0, 1+ and 2+ cases) and “CD44_high” (3+ cases). GC patients whose tumors were classified as “CD44_high” had significantly worse OS than those with absent/low CD44v6 expression (median OS ∼ 15 months vs 51 months, *P* < .0001; Fig 1C). When comparing the clinical-pathological features of patients bearing CD44_absent/low with those bearing CD44_high tumors, we found that CD44_high patients more often presented vascular invasion (*P* < .005), tumor cells in the surgical margins (*P* < .01) and perineural invasion (*P* < .05) (Table 1). Multivariate analysis further demonstrated that CD44_high was an independent factor of poor prognosis in this series (Fig 1D). Additionally, and somehow expected, high pTNM stage (*P* < .0001), older age at onset (*P* <.0001) and presence of tumor cells in surgical margins (*P* = .003) were also identified as independent poor prognosis factors in this series.

**Table 1:** Clinical-pathological associations with extent of CD44v6 expression (absent/low *vs*. high) in gastric tumors.

| Variables | Total No. patients (n= 334) | CD44v6 absent/low (n= 287) | CD44v6 high (n= 47) | P-value |
| --- | --- | --- | --- | --- |
| **Age (years)** |  |  |  | > .05 |
| Mean | 67.6 | 67.3 | 69.3 |  |
| Standard deviation | 11.9 | 11.9 | 11.9 |  |
| **Gender** |  |  |  | > .05 |
| Male | 188 (56.3%) | 167 (58.2%) | 21 (44.7%) |  |
| Female | 146 (43.7%) | 120 (41.8%) | 26 (55.3%) |  |
| Male:Female ratio | 1.3:1 | 1.4:1 | 0.8:1 |  |
| **Lauren classification** |  |  |  | > .05 |
| Intestinal | 156 (46.7%) | 139 (48.4%) | 17 (36.2%) |  |
| Diffuse | 44 (13.2%) | 39 (13.6%) | 5 (10.6%) |  |
| Mixed | 86 (25.7%) | 68 (23.7%) | 18 (38.3%) |  |
| Indeterminate | 48 (14.4%) | 41 (14.3%) | 7 (14.9%) |  |
| **WHO classification** |  |  |  | > .05 |
| Tubular | 146 (43.7%) | 129 (44.9%) | 17 (36.2%) |  |
| Papillary | 2 (0.6%) | 2 (0.7%) | 0 (0.0%) |  |
| Mucinous | 7 (2.1%) | 7 (2.4%) | 0 (0.0%) |  |
| Poorly cohesive | 38 (11.4%) | 32 (11.1%) | 6 (12.8%) |  |
| Other types | 141 (42.2%) | 117 (40.8%) | 24 (51.1%) |  |
| **Growth pattern** |  |  |  | > .05 |
| Expansive | 61 (18.3%) | 55 (19.2%) | 6 (12.8%) |  |
| Infiltrative | 259 (77.5%) | 220 (76.7%) | 39 (83.0%) |  |
| Unclassified | 14 (4.2%) | 12 (4.2%) | 2 (4.3%) |  |
| **Wall invasion** |  |  |  | > .05 |
| Mucosa + Submucosa | 79 (23.7%) | 74 (25.8%) | 5 (10.6%) |  |
| Muscular | 42 (12.6%) | 36 (12.5%) | 6 (12.8%) |  |
| Subserosa + Serosa | 201 (60.2%) | 167 (58.2%) | 34 (72.3%) |  |
| Other organs | 12 (3.6%) | 10 (3.5%) | 2 (4.3%) |  |
| **Lymphatic permeation** | 332*1 |  |  | > .05 |
| Absent | 103 (31.0%) | 94 (33.0%) | 9 (19.1%) |  |
| Present | 229 (69.0%) | 191 (67.0%) | 38 (80.9%) |  |
| **Perineural invasion** | 333*1 |  |  | .043 |
| Absent | 173 (52.0%) | 155 (54.2%) | 18 (38.3%) |  |
| Present | 160 (48.0%) | 131 (45.8%) | 29 (61.7%) |  |
| **Vascular invasion** | 331*1 |  |  | .002 |
| Absent | 138 (41.7%) | 128 (45.1%) | 10 (21.3%) |  |
| Present | 193 (58.3%) | 156 (54.9%) | 37 (78.7%) |  |
| **Surgical margins** | 333*1 |  |  | .009 |
| R0 | 298 (89.5%) | 261 (91.3%) | 37 (78.7%) |  |
| R1/R2 | 35 (10.5%) | 25 (8.7%) | 10 (21.3%) |  |
| **Depth of invasion (T)** |  |  |  | > .05 |
| pT1 | 79 (23.7%) | 74 (25.8%) | 5 (10.6%) |  |
| pT2 | 79 (23.7%) | 68 (23.7%) | 11 (23.4%) |  |
| pT3-T4 | 176 (52.7%) | 145 (50.5%) | 31 (66.0%) |  |
| **Lymph node metastases (N)** | 333*1 |  |  | > .05 |
| Absent (pN0) | 129 (38.7%) | 116 (40.6%) | 13 (27.7%) |  |
| Present (pN+) | 204 (61.3%) | 170 (59.4%) | 34 (72.3%) |  |
| **Distant metastases (M)** |  |  |  | > .05 |
| Absent | 265 (79.3%) | 229 (79.8%) | 36 (76.6%) |  |
| Present | 69 (20.7%) | 58 (20.2%) | 11 (23.4%) |  |
| **pTNM Staging** |  |  |  | > .05 |
| I | 109 (32.6%) | 100 (34.8%) | 9 (19.1%) |  |
| II | 87 (26.0%) | 72 (25.1%) | 15 (31.9%) |  |
| III | 69 (20.7%) | 57 (19.9%) | 12 (25.5%) |  |
| IV | 69 (20.7%) | 58 (20.2%) | 11 (23.4%) |  |
| **Disease Recurrence** | 265*2 |  |  | > .05 |
| Absent | 192 (72.5%) | 168 (73.4%) | 24 (66.6%) |  |
| Present | 73 (27.5%) | 61 (26.6%) | 12 (33.3%) |  |
| **Treatment Regimen** | 326*1 |  |  | > .05 |
| Surgery alone | 203 (62.3%) | 175 (62.7%) | 28 (59.6%) |  |
| Pt-based therapy | 113 (34.7%) | 94 (33.7%) | 19 (40.4%) |  |
| Non-Pt based therapy | 10 (3.1%) | 10 (3.6%) | 0 (0.0%) |  |

\*<sup>1</sup> Remaining data not available; \*<sup>2</sup> Stage IV patients were excluded. pTNM (pathological tumor-node-metastasis).

Remaining data not available; *2 Stage IV patients were excluded. pTNM (pathologica tumor-node-metastasis).

In summary, we identified a clear cut-off for defining overexpression of CD44v6 in GC (>50% tumor cells overexpressing membranous CD44v6), which demonstrates prognostic value, likely due to greater ability of CD44_high cells to access the vasculature and perineural space, and to spread throughout the stomach wall.

### Adding chemotherapy to surgery benefits particularly patients presenting CD44_high tumors

Given that patients bearing “CD44_high” tumors present worse prognosis, we hypothesize that they may also respond worse to cisplatin-based therapy. Indeed, we observed that “CD44v6-high” patients have worse median OS regardless of receiving chemotherapy in addition to surgery or not (Figs 2A and B). However, administration of chemotherapy to CD44_high patients resulted in a 6-fold increase in their median OS, from 4 to 24 months (Figs 2A and B). Although not as striking, the median OS of patients with CD44_absent/low tumors almost doubled (from 20 to 38 months) when treated with conventional chemotherapy in addition to surgery (Figs 2A and B). This benefit was mainly associated with patients with CD44_2+ (Fig 2E). Overall, these data support chemotherapy not only in CD44v6-high (CD44_3+) patients, but also in CD44_2+ patients, and fails to support our initial hypothesis that the poor prognosis of CD44v6-high patients is related to weak response to chemotherapy. Indeed, we further verified that the greater the percentage of tumor cells expressing CD44v6, the greater the treatment benefit reflected in the median OS (Figs 2C-G). For instance, CD44_1+, 2+ and 3+ patients had a 2.6 (not statistically significant), 3.8 and 6-fold improvement in OS when treated with chemotherapy in addition to surgery, respectively (*P* = .004, Figs 2D-F). In contrast, no OS improvement was observed when administering chemotherapy to CD44v6 negative patients (Fig 2C).

Although the extent of CD44v6 expression does not seem to affect RFS (Appendix Figs 4A-G), the mean time to relapse decreases with increasing percentage of CD44v6+ cells in the tumor (∼28, 21 and 15 months for CD44v6 1+, 2+ and 3+, respectively; Appendix Fig 4H) in patients treated with surgery and chemotherapy. This is true after chemotherapy, when considering only the sub-set of patients that relapsed. These results suggest that CD44v6+ cells in the tumor are likely involved in triggering a faster tumor relapse following chemotherapy. In patients from the CD44_0 group, chemotherapy seems to decrease time to relapse as compared with surgery alone (from 19 to 13 months), unlike what occurs in the CD44v6+ groups (Appendix Fig 4H). These data raise the importance of considering the CD44v6 sub-categorization when selecting patients for chemotherapy, as: (1) chemotherapy significantly improved OS and time to relapse specifically in CD44_2+ and 3+ patients, and; (2) chemotherapy does not influence OS and decreases time to relapse in CD44v6 negative (CD44_0) patients.

### CD44v6 overexpression in GC cell lines influences response to cisplatin treatment

We next wanted to understand how does CD44v6 overexpression influences response to conventional chemotherapeutic agents used in GC treatment, namely cisplatin and 5-fluorouracil (5-FU). For this, we generated three isogenic GC cell lines: mock cells lacking CD44, CD44s and CD44v6 expressing cells (Appendix Fig 5).

While no differences were observed between the isogenic cells for doubling-time, invasion, cell-adhesion and motility, tumorigenic potential *in vivo* (Appendix Fig 6); CD44v6 overexpressing cells survived better to cisplatin treatment, by decreasing apoptosis levels (Fig 3A and C), but not to 5-FU treatment (Fig 3B). To validate these findings, we depleted CD44v6 expression from cells endogenously expressing CD44v6 (Appendix Fig 7). Indeed, CD44v6 depletion increased cisplatin-induced apoptosis (Fig 3D), validating our results. Since some CD44v6 effectors are also implicated in cisplatin resistance, namely STAT3 and P38, we evaluated their modulation in our experimental models. STAT3 was rapidly activated in CD44v6 overexpressing cells in response to cisplatin (Appendix Fig 8). Although this might explain the increased cell survival in CD44v6 overexpressing cells, this behavior was also observed in CD44s cells. This data was not further clarified neither when analyzing P38 expression in the isogenic model nor in cell lines endogenously expressing CD44v6 (data not shown).

**Fig 3:**
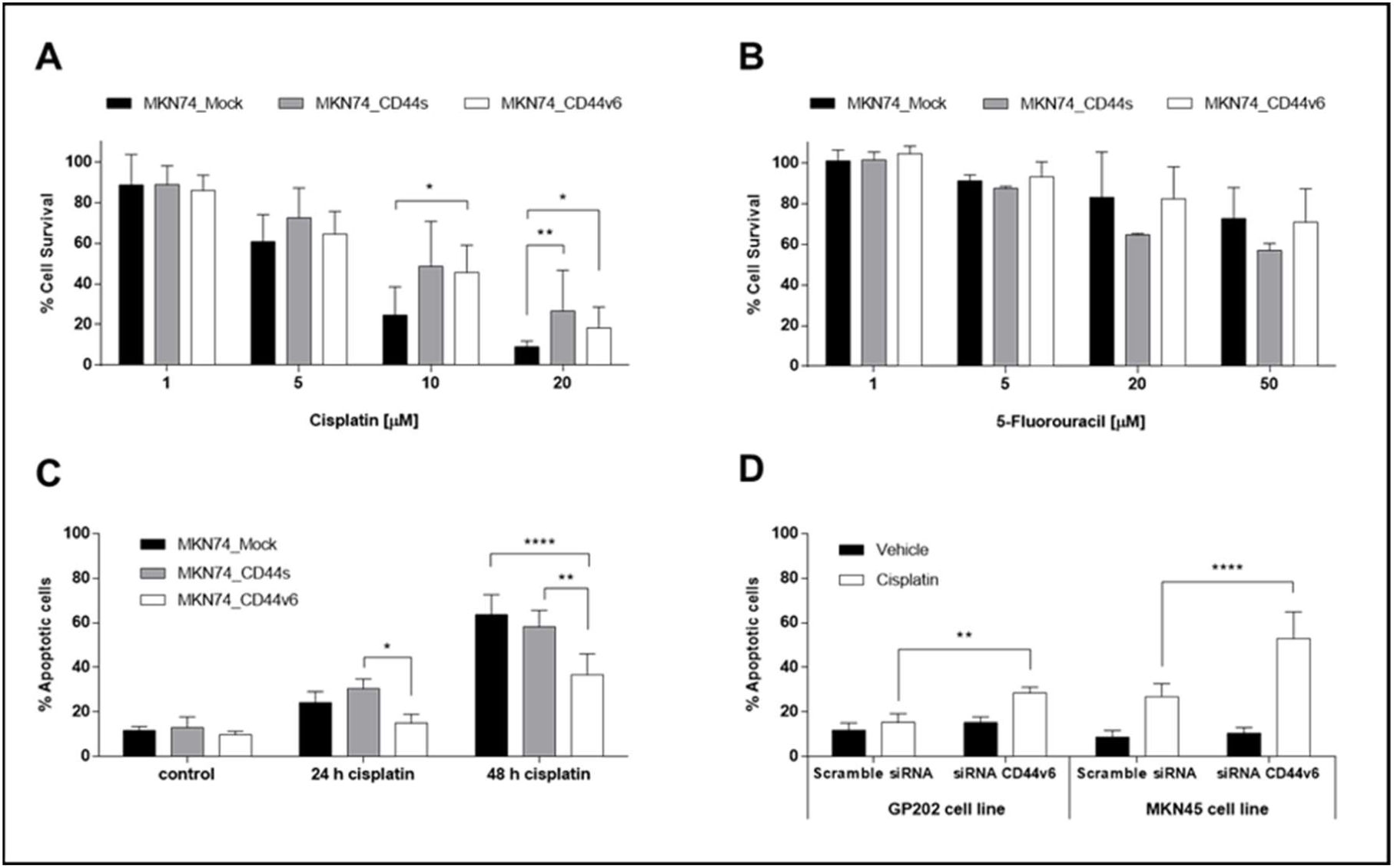
Assessing the response of the isogenic MKN74 cells to conventionally used chemotherapeutic agents: **(A)** % Cell survival upon incubation with cisplatin or **(B)** 5-FU for 48 h (compared to vehicle control) in MKN74 cells; **(C)** % Apoptotic cells in MKN74 cells incubated with 10 µM cisplatin or vehicle (0.9% NaCl) for 48 h; **(D)** % Apoptotic cells in GP202 and MKN45 cell lines in response to 48 h treatment with cisplatin or vehicle, following a 24 h incubation with CD44v6 or scramble siRNAs. The cisplatin concentrations used are considered to be clinically relevant. Results are expressed as the average + SD of at least three independent experiments. Statistically significant results was determined by Two-way ANOVA with Tukey’s multiple comparisons test (* *P* <.05; ** *P* < .001; **** *P* < .0001).

Altogether, these *in vitro* studies clearly and consistently indicated that CD44v6 overexpression increases survival of GC cells in response to cisplatin treatment.

### CD44v6 expressing cells may contribute to patient relapse after chemotherapy

We observed that patients’ tumors with greater CD44v6+ cell populations relapse faster, following chemotherapy than those with small fractions of this sub-population (Appendix Fig 4H). In addition, our *in vitro* results show that CD44v6+ cells survive better than CD44v6-cells in response to cisplatin (included in the chemotherapeutic regimen of most patients studied; Appendix Table 2). So, we hypothesized that upon chemotherapy, CD44v6+ cells prevail over the CD44v6-counterpart, triggering faster tumor relapse and dominating recurring lesions. We therefore analyzed the CD44v6 *status* in surgically-resected tumors from patients treated with neo-adjuvant chemotherapy or chemo-naïve. Supporting our hypothesis, 50% of tumors treated prior to surgery were CD44_3+, comparing to 25% of chemo-naïve tumors (Fig 4A). Despite the low number of patients, these results suggest that chemotherapy may increase the abundance of CD44v6+ cell population in gastric cancers. We tested this hypothesis *in vitro* by incubating co-cultures of CD44v6+ and CD44v6-cells with cisplatin and subsequently allowing them to recover for 15 days (as an attempt to mimic what happens in patients that receive chemotherapy and then present tumor relapse). We showed that upon recovery, CD44v6+ cells represent >80% of the cell culture, suggesting that CD44v6 is involved in GC cell overgrowth after cisplatin treatment (Figs 4B and C).

**Fig 4:**
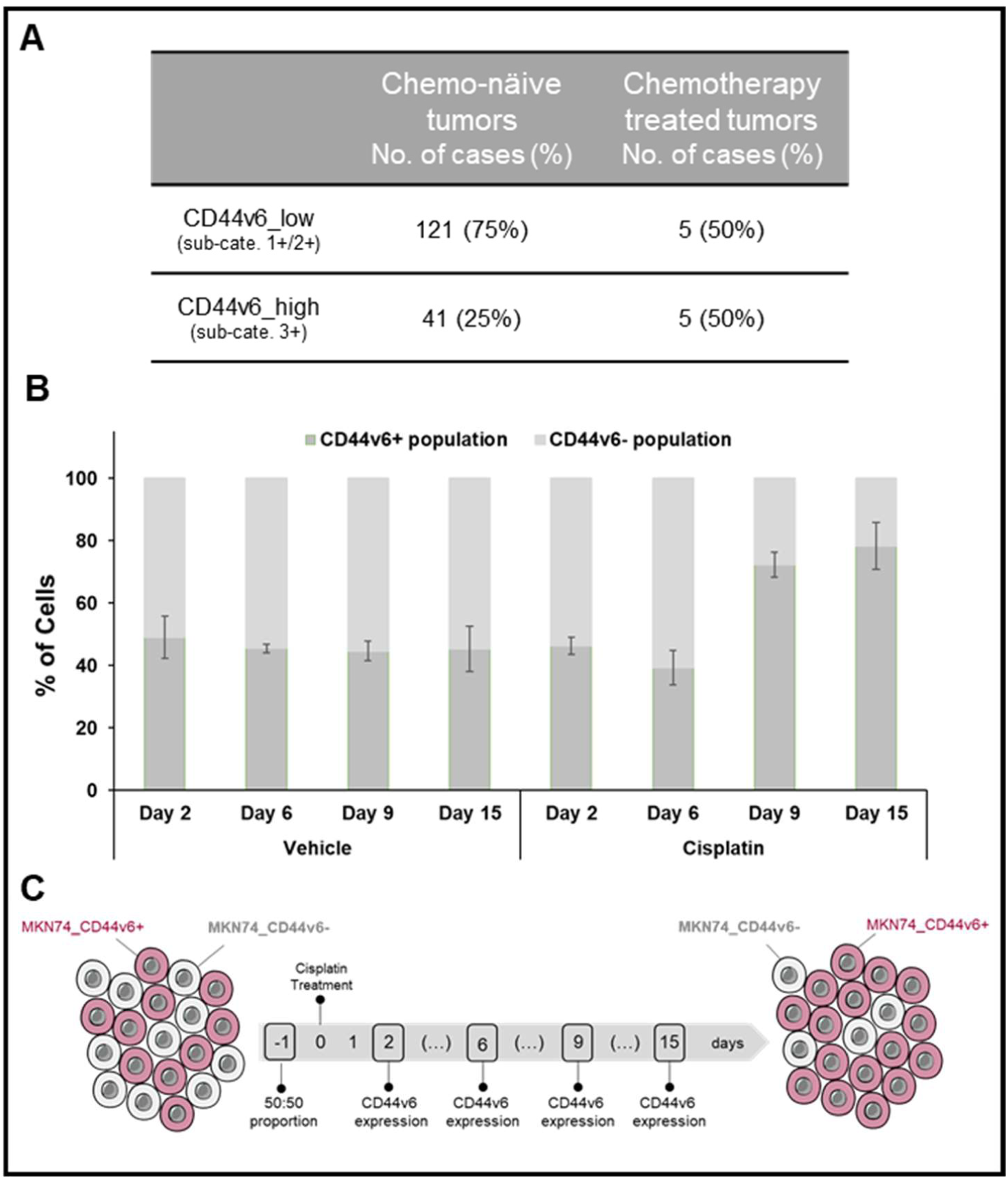
In heterogeneous tumors, CD44v6+ cells may be enriched following treatment with chemotherapeutic agents **(A)** frequency of CD44_low (comprising CD44v6 sub-categories 1+ and 2+) and CD44_high in chemo-naïve (comprising both tumors from patients that did not receive chemotherapy in addition to surgery and also patients that only received chemotherapy following tumor resection); **(B)** % of CD44v6+ cells in co-cultures of CD44v6+ and CD44v6-MKN74 cells (MKN74_CD44v6 and MKN74_Mock cells, respectively) following a 6 hour incubation with cisplatin (or vehicle control) and up to 15 days of recovery. At the end of the experiment > 80% of the cell culture is composed of CD44v6+ cells; **(C)** Scheme depicting the cell enrichment experiment.

## Discussion

Herein, we established a protocol to evaluate CD44v6 in GC, according to the percentage of tumor cells expressing membranous CD44v6. The categorization of CD44v6 expression was further refined through correlation with OS of GC patients, and allowed demonstrating that in GC, CD44v6 is an independent marker of poor prognosis with further potential as a predictive marker of response to conventional chemotherapy. Our data also shed light into GC molecular heterogeneity and its relation with therapy, by suggesting that the CD44v6+ population may be driving recurrence following conventional chemotherapy, in tumors composed of CD44v6+ and CD44v6-cell populations. Approximately 14% of GC patients present with CD44_high tumors (Fig 5A), which have a significantly worse median OS compared with CD44_absent/low patients. Importantly, this occurs independently of TNM staging (Appendix Fig 9A-D), as demonstrated by multivariate analysis. Studies so far reporting an association between CD44v6 expression and worse GC patient survival,^18,19^enclosed low patients’ number and failed to demonstrate prognostic value for CD44v6 expression independently of TNM staging. Our study addresses this issue in the largest GC patient cohort tested so far, and demonstrates that CD44v6 is truly an independent marker of poor prognosis, as it has been described for colorectal cancer.^25^

**Fig 5:**
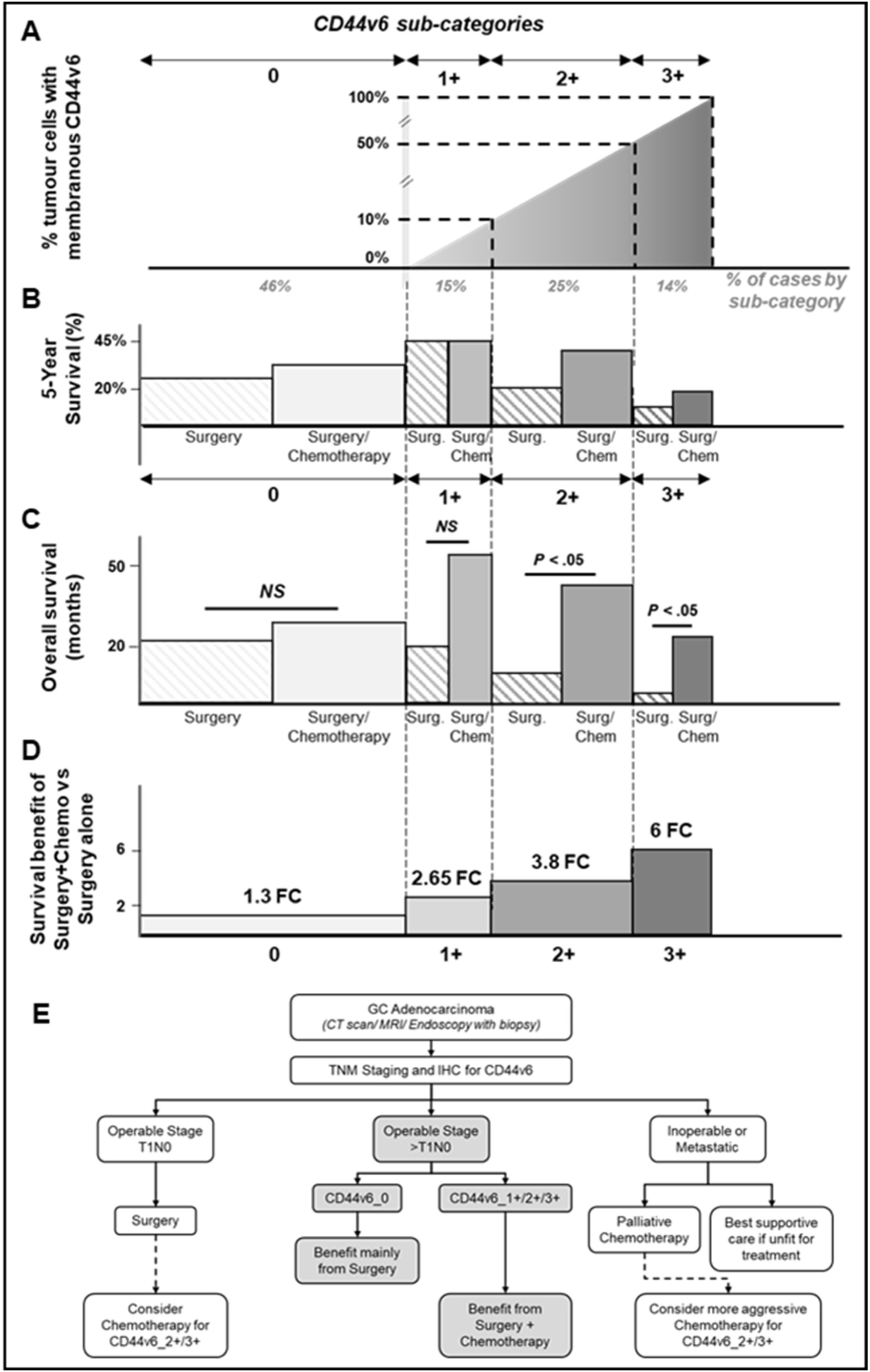
(A) Classification of GC in sub-categories of CD44v6 expression and % of cases per sub-category; **(B)** 5-year survival of patients treated with surgery alone and with surgery plus chemotherapy, per CD44v6 sub-category; **(C)** median OS of patients treated with surgery alone and with surgery plus chemotherapy, per CD44v6 sub-category; **(D)** GC patient survival benefit of adding chemotherapy to surgery, expressed as fold change (FC) between the OS of patients treated with surgery + chemotherapy with the OS of patients treated with surgery alone; **(E)** proposed scheme for improving GC patient stratification using CD44v6 as a marker for treatment selection (adapted from ^28^). The obtained data show that patients with operable >T1N0 and CD44v6+ tumors benefit from receiving chemotherapy in addition to surgery. In addition, our data suggests patients with T1N0 and CD44v6+ tumors may benefit from additional chemotherapy, and that patients with inoperable or metastatic GC with CD44v6+ tumors may benefit from more aggressive chemotherapeutic strategies, highlighting the importance of performing additional studies on this issue.

We further demonstrated that CD44v6 expression in tumors predicts benefit from application of conventional chemotherapy in addition to surgery. This benefit is directly proportional to the fraction of CD44v6+ cells in tumors, with patients whose tumors lack CD44v6 expression not benefiting from chemotherapy treatment (Fig 5B-5D). The latter also appear to have worse RFS and to relapse sooner than CD44_0 patients not receiving chemotherapy in addition to surgery. To the best of our knowledge, this is the first report demonstrating CD44v6 as a useful marker to predict therapy response in GC, similarly to what was shown in colorectal cancer, where patients with moderate or strong CD44v6 tumor expression respond better to chemotherapy with irinotecan.^26^

Nearly half of the tumors herein studied possess some degree of heterogeneity regarding CD44v6 expression (with CD44v6+ and CD44v6-cell populations co-existing in the same tumor), which may have important consequences in terms of clinical outcome. Our *in vitro* data shows that CD44v6 expression promotes survival of GC cells after cisplatin treatment, as others have shown for several tumor models,^15,23,24^and in co-cultures of CD44v6+ and CD44v6– cells. Our patient data also suggests enrichment in the CD44v6+ cell population after neo-adjuvant treatment and in relapses, but validation in larger and/or independent GC patient series is still needed.

CD44/CD44v are considered cancer stem cell markers in several tumor types, including GC,^13,14^supporting our hypothesis that CD44v6+ tumor cells may be potentiating relapse. If our findings are confirmed, strategies to specifically eliminate CD44v6+ cells could lead to decreased recurrence and improved patient survival. One possible strategy is the targeting of CD44v6 downstream effectors, as direct targeting of CD44v6 ought to be avoided due to the lethal skin toxicity side effects described in a Phase I clinical trial using a highly potent antimicrotubule agent coupled to a monoclonal antibody against CD44v6.^27^TNM staging has long been the most important tool to assess prognosis in GC. Our data shows that evaluating CD44v6 by IHC may provide oncologists with additional and important information for stratifying GC patient for treatment. Namely, if CD44v6 IHC is performed in GC biopsies and turns out CD44_high, this information may be used to recommend surgery combined with chemotherapy. This would be particularly important for patients with CD44v6 2+ or 3+ tumors diagnosed in stage I and stage II, who often do not undergo chemotherapy. Moreover, finding tumors lacking CD44v6 expression could help oncologists selecting patients less likely to benefit from chemotherapy, hence avoiding its related toxicity side effects (Fig 5E).

## Conclusion

Our study is the first one to identify CD44v6 as a potential predictive marker of response to conventional chemotherapy. In addition, we provide compelling evidence that consolidates CD44v6 is an independent marker of poor prognosis, and a likely marker of faster relapse. Importantly, our data supports selection of patients with high CD44v6 expressing tumors for conventional chemotherapy in addition to surgery, even in TNM stage I and stage II patients, who are generally treated with surgery alone.

## Supporting information

## Acknowledgements

This work was supported by FEDER - Fundo Europeu de Desenvolvimento Regional funds through the COMPETE 2020 – Operacional Programme for Competitiveness and Internationalisation (POCI), Portugal 2020, and by Portuguese funds through FCT – Fundação para a Ciência e a Tecnologia/Ministério da Ciência, Tecnologia e Inovação in the framework of the project “Institute for Research and Innovation in Health Sciences” (POCI-01-0145-FEDER-007274). This work was also financed by the projects NORTE-01-0145-FEDER-000003 (DOCnet) and NORTE-07-0124-FEDER-000029 - supported by Norte Portugal Regional Programme (NORTE 2020), under the PORTUGAL 2020 Partnership Agreement, through the European Regional Development Fund (ERDF) - and project PTDC/CTM-NAN/120958/2010, from FCT.

Carla Pereira was supported by the grant SFRH/BD/113031/2015 and Daniel Ferreira by the grant PD/BD/105976/2014.

Gabriela M. Almeida was supported by the Investigator FCT Program 2013 (IF/00615/2013), POPH - QREN Type 4.2, European Social Fund and Portuguese Ministry of Science and Technology (MCTES).

## References

[1] Bray F, Ferlay J, Soerjomataram I, et al. Global cancer statistics 2018: GLOBOCAN estimates of incidence and mortality worldwide for 36 cancers in 185 countries. CA Cancer J Clin 10.3322/caac.21492. [Epub ahead of print], 2018

[2] Oba K, Paoletti X, Bang YJ, et al: Role of chemotherapy for advanced/recurrent gastric cancer: an individual-patient-data meta-analysis. Eur J Cancer 49:1565-1577, 2013

[3] Cervantes A, Roda D, Tarazona N, et al: Current questions for the treatment of advanced gastric cancer. Cancer Treat Rev 39:60-67, 2013

[4] Durães C, Almeida GM, Seruca R, et al: Biomarkers for gastric cancer: prognostic, predictive or targets of therapy? Virchows Arch 464:367-378, 2014

[5] Bang YJ, Van Cutsem E, Feyereislova A, et al: Trastuzumab in combination with chemotherapy versus chemotherapy alone for treatment of HER2-positive advanced gastric or gastro-oesophageal junction cancer (ToGA): a phase 3, open-label, randomised controlled trial. Lancet 376:687-697, 2010

[6] Fuchs CS, Tomasek J, Yong CJ, et al: Ramucirumab monotherapy for previously treated advanced gastric or gastro-oesophageal junction adenocarcinoma (REGARD): an international, randomised, multicentre, placebo-controlled, phase 3 trial. Lancet 383:31-39, 2014

[7] Ponta H, Wainwright D, Herrlich P: The CD44 protein family. Int J Biochem Cell Biol 30: 299–305, 1998

[8] Ponta H, Sherman L, Herrlich PA: CD44: from adhesion molecules to signalling regulators. Nat Rev Mol Cell Biol 4: 33–45, 2003

[9] Prochazka L, Tesarik R, Turanek J: Regulation of alternative splicing of CD44 in cancer. Cell Signal 26:2234–2239, 2014

[10] Ruiz P, Schwarzler C, Gunthert U: CD44 Isoforms during Differentiation and Development. Bioessays 17:17–24, 1995

[11] Naor D, Wallach-Dayan SB, Zahalka MA, et al: Involvement of CD44, a molecule with a thousand faces, in cancer dissemination. Seminars in Cancer Biology 18: 260–267, 2008

[12] Senbanjo LT, Chellaiah MA: CD44: A Multifunctional Cell Surface Adhesion Receptor Is a Regulator of Progression and Metastasis of Cancer Cells. Front Cell Dev Biol 5:18, 2017

[13] Yan Y, Zuo X, Wei D: Concise Review: Emerging Role of CD44 in Cancer Stem Cells: A Promising Biomarker and Therapeutic Target. Stem Cells Transl Med 4:1033–1043, 2015

[14] Todaro M, Gaggianesi M, Catalano V, et al: CD44v6 is a marker of constitutive and reprogrammed cancer stem cells driving colon cancer metastasis. Cell Stem Cell 14:342–356, 2014

[15] Zöller M: CD44, Hyaluronan, the Hematopoietic Stem Cell, and Leukemia-Initiating Cells. Front Immunol 6:235, 2015

[16] da Cunha CB, Oliveira C, Wen X, et al: De novo expression of CD44 variants in sporadic and hereditary gastric cancer. Lab Invest 90:1604–1614, 2010

[17] Chen Y, Fu Z, Xu S, et al: The prognostic value of CD44 expression in gastric cancer: a meta-analysis. Biomed Pharmacother 68:693–697, 2014

[18] Xie JW, Chen PC, Zheng CH, et al. Evaluation of the prognostic value and functional roles of CD44v6 in gastric cancer. J Cancer Res Clin Oncol 141:1809–1817, 2015

[19] Xu YY, Guo M, Yang LQ, et al: Regulation of CD44v6 expression in gastric carcinoma by the IL-6/STAT3 signaling pathway and its clinical significance. Oncotarget 8:45848–45861, 2017

[20] Rodrigues M, Vitó I, Santos R, et al: Establishment of a Tumour Bank: the experience of the Department of Pathology of Hospital S. João (Porto, Portugal). Cell Tissue Bank 10:75–77, 2009

[21] Gärtner F, David L, Seruca R, et al: Establishment and characterization of two cell lines derived from human diffuse gastric carcinomas xenografted in nude mice. Virchows Arch 428:91–98, 1996

[22] Bordeira-Carrico R, Ferreira D, Mateus DD, et al: Rescue of wild-type E-cadherin expression from nonsense-mutated cancer cells by a suppressor-tRNA. Eur J Hum Genet 22:1085–1092, 2014

[23] Lv L, Liu HG, Dong SY, et al: Upregulation of CD44v6 contributes to acquired chemoresistance via the modulation of autophagy in colon cancer SW480 cells. Tumour Biol 37:8811–8824, 2016

[24] Mooney KL, Choy W, Sidhu S, et al: The role of CD44 in glioblastoma multiforme. J Clin Neurosci 34:1–5, 2016

[25] Wang JL, Su WY, Lin YW, et al: CD44v6 overexpression related to metastasis and poor prognosis of colorectal cancer: A meta-analysis. Oncotarget 8:12866–12876, 2017

[26] Bendardaf R, Lamlum H, Ristamäki R, et al: CD44 variant 6 expression predicts response to treatment in advanced colorectal cancer. Oncol Rep 11:41– 45, 2004

[27] Riechelmann H, Sauter A, Golze W, et al: Phase I trial with the CD44v6-targeting immunoconjugate bivatuzumab mertansine in head and neck squamous cell carcinoma. Oral Oncol 44:823–829, 2008

[28] Lordick F, Allum W, Carneiro F, et al: Unmet needs and challenges in gastric cancer: the way forward. Cancer Treat Rev 40:692–700, 2014

