## Supplementary material for "CD44v6 expression is a novel predictive marker of therapy response and poor prognosis in gastric cancer patients"

### *Supplemental Figures*

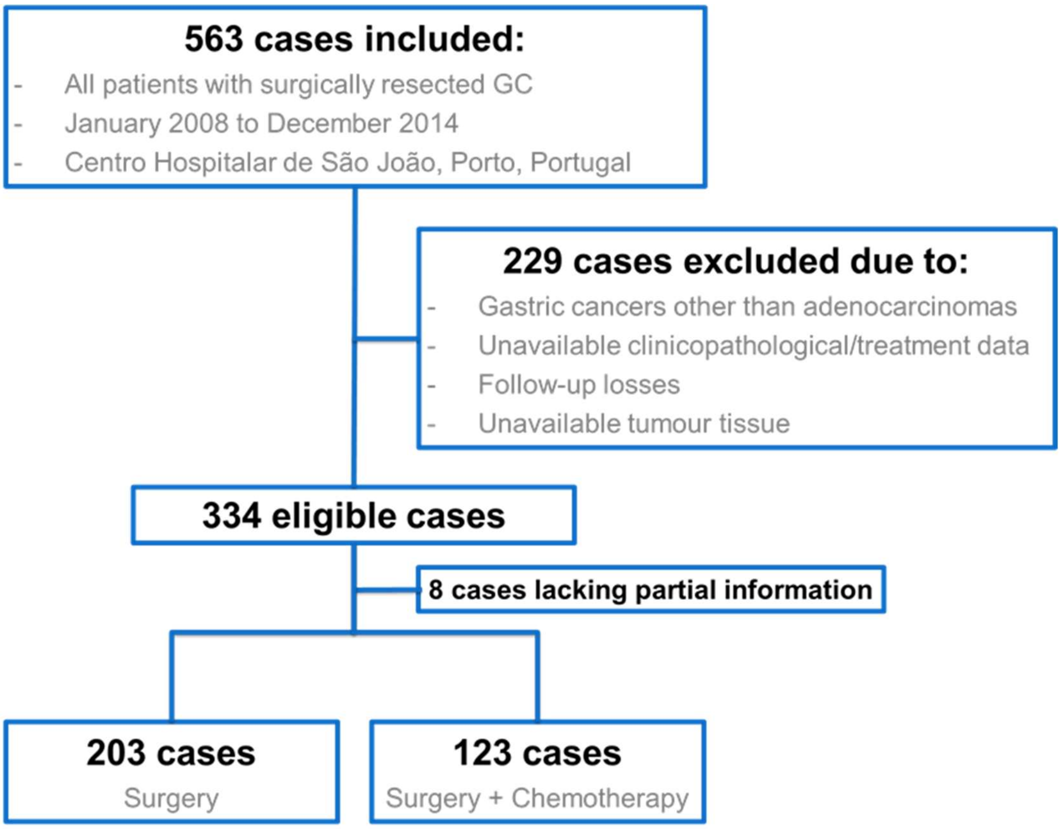

**Appendix Fig 1:** GC cohort profile. From the 334 gastric adenocarcinoma patients, eligible for this study, 203 were treated only with surgery and 123 patients were treated with surgery plus chemotherapy.

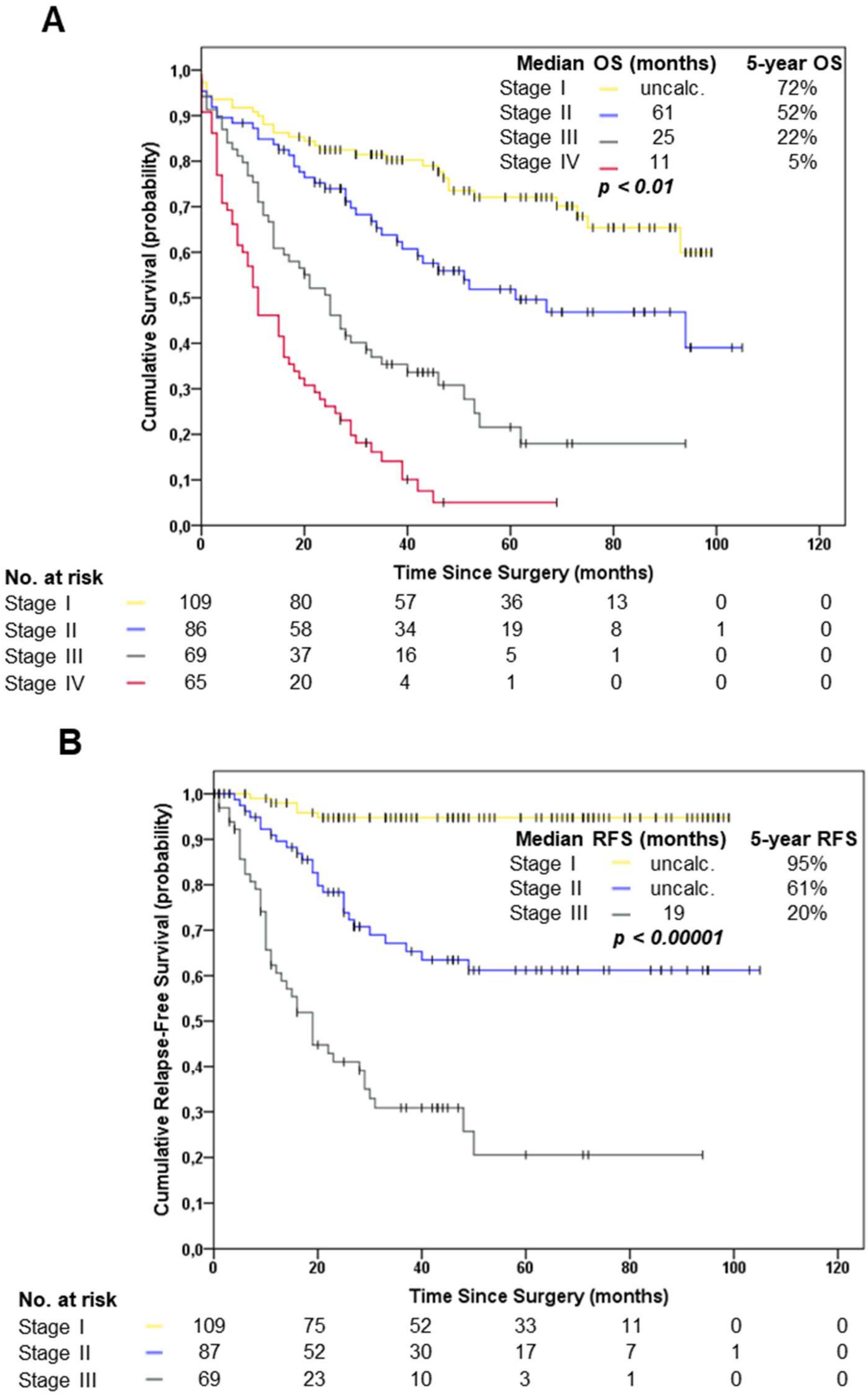

**Appendix Fig 2: (A)** Kaplan-Meier estimates showing OS and **(B)** RFS of GC patients in the cohort, according to pTNM staging.

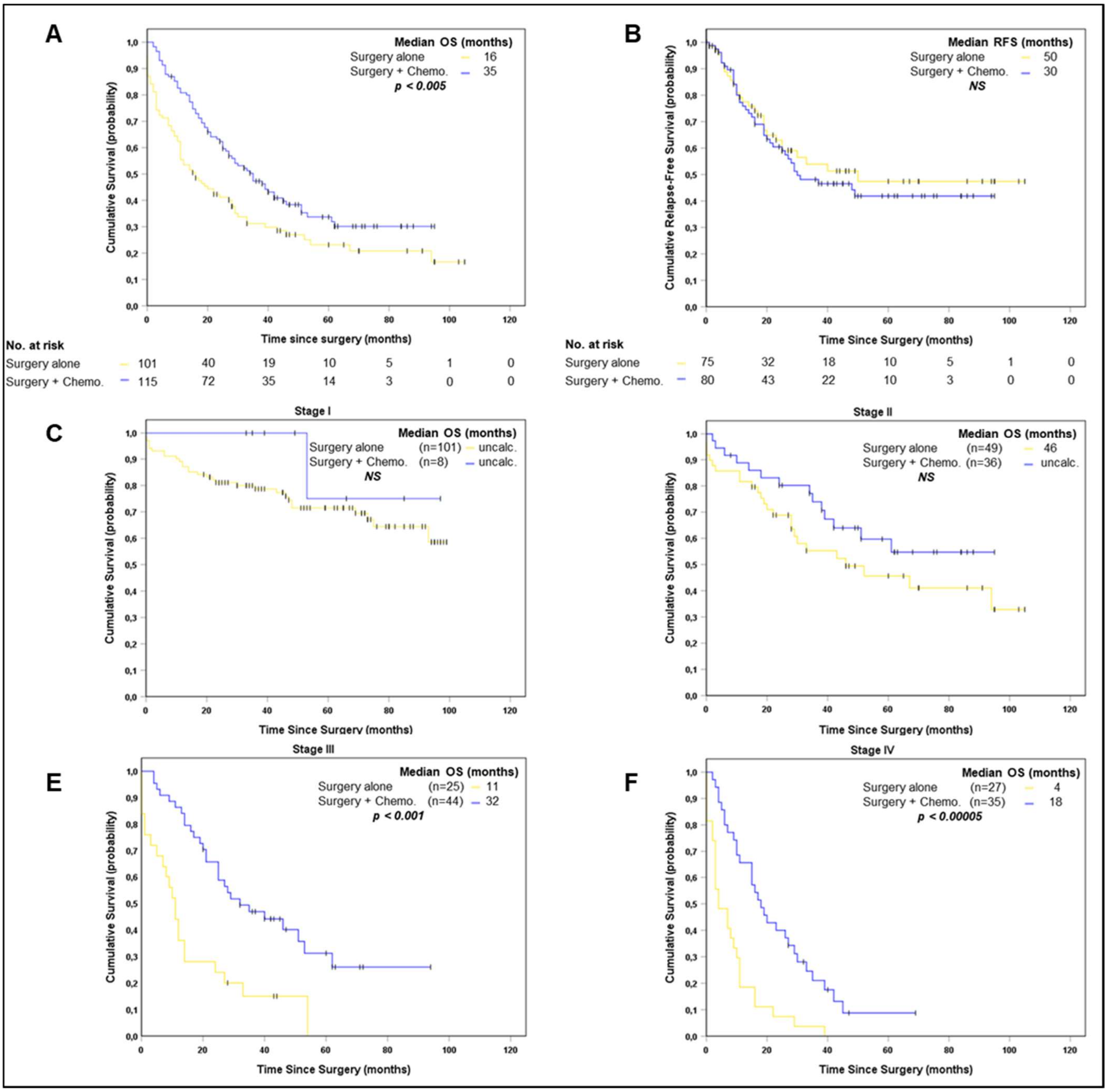

**Appendix Fig 3: (A)** Kaplan-Meier estimates showing OS and **(B)** RFS of GC patients in the cohort, according to whether they were treated with surgery alone or with surgery + chemotherapy. Since patients that received chemotherapy in addition to surgery were mostly stage II to IV patients, when evaluating the benefit of chemotherapy on OS, pTNM stage I patients were excluded from the analysis as the majority of these patients (> 90%) had surgery alone, and including them in this analysis would introduce a bias in results; **(C)** Kaplan-Meier estimates showing OS in GC patients from TNM Stage I, **(D)** Stage II, **(E)** Stage III and **(F)** Stage IV, according to the therapy they received.

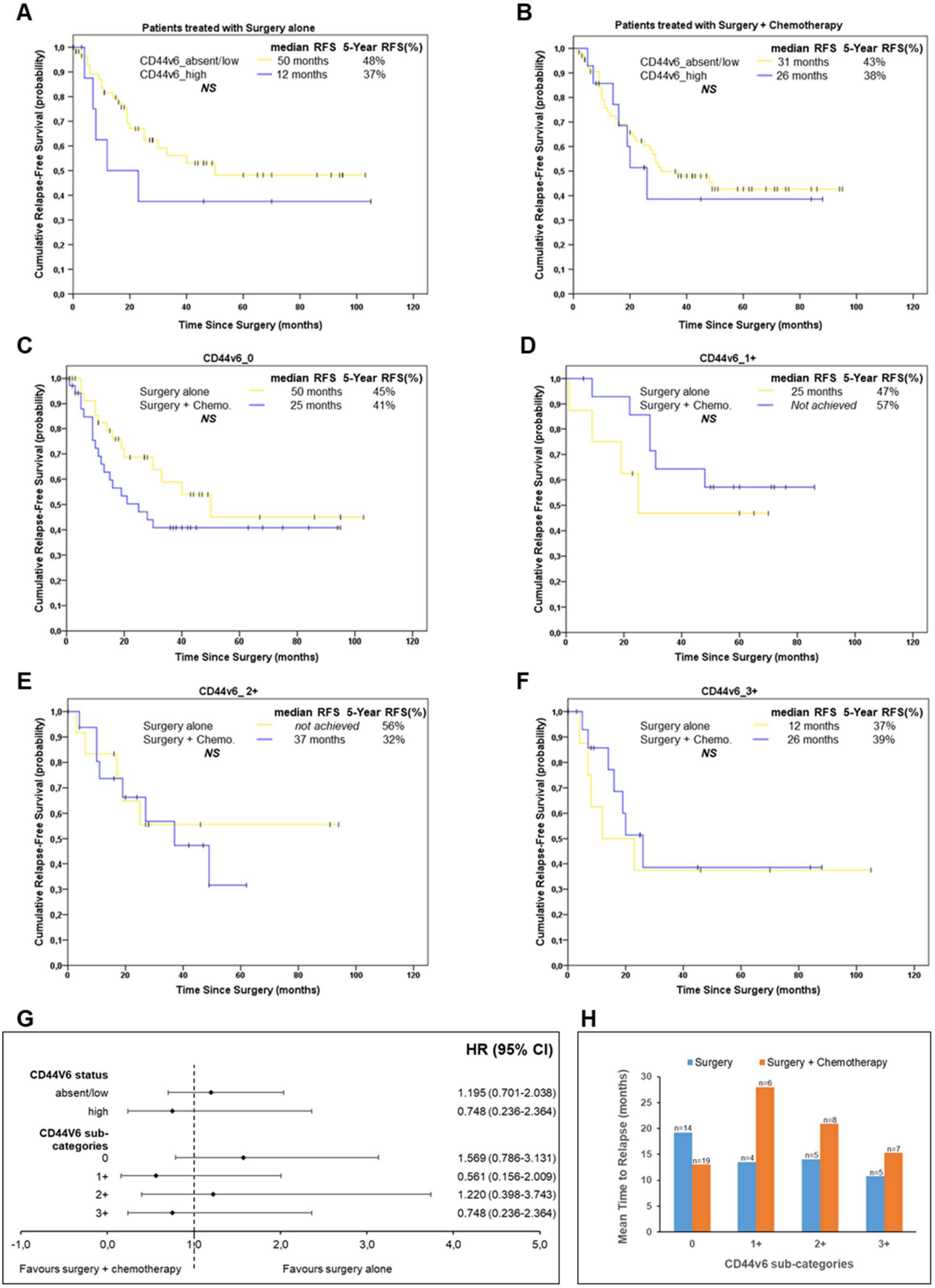
A
Patients treated with Surgery alone
median RFS 5-Year RFS(%)
CD44v6_absent/low 50 months 48%
CD44v6_high 12 months 37%
NS
Cumulative Relapse-Free Survival (probability)
Time Since Surgery (months)
B
Patients treated with Surgery + Chemotherapy
median RFS 5-Year RFS(%)
CD44v6_absent/low 31 months 43%
CD44v6_high 26 months 38%
NS
Cumulative Relapse-Free Survival (probability)
Time Since Surgery (months)
C
CD44v6_0
median RFS 5-Year RFS(%)
Surgery alone 50 months 45%
Surgery + Chemo. 25 months 41%
NS
Cumulative Relapse-Free Survival (probability)
Time Since Surgery (months)
D
CD44v6_1+
median RFS 5-Year RFS(%)
Surgery alone 25 months 47%
Surgery + Chemo. Not achieved 57%
NS
Cumulative Relapse Free Survival (probability)
Time Since Surgery (months)
E
CD44v6_2+
median RFS 5-Year RFS(%)
Surgery alone not achieved 56%
Surgery + Chemo. 37 months 32%
NS
Cumulative Relapse-Free Survival (probability)
Time Since Surgery (months)
F
CD44v6_3+
median RFS 5-Year RFS(%)
Surgery alone 12 months 37%
Surgery + Chemo. 26 months 39%
NS
Cumulative Relapse-Free Survival (probability)
Time Since Surgery (months)
G
HR (95% CI)
CD44V6 status
absent/low 1.195 (0.701-2.038)
high 0.748 (0.236-2.364)
CD44V6 sub-categories
0 1.569 (0.786-3.131)
1+ 0.561 (0.156-2.009)
2+ 1.220 (0.398-3.743)
3+ 0.748 (0.236-2.364)
-1,0 0,0 1,0 2,0 3,0 4,0 5,0
Favours surgery + chemotherapy
Favours surgery alone
H
Surgery
Surgery + Chemotherapy
Mean Time to Relapse (months)
n=14 n=19 n=4 n=6 n=5 n=8 n=5 n=7
0 1+ 2+ 3+
CD44v6 sub-categories

**Appendix Fig 4: (A)** Kaplan-Meier estimates showing RFS of patients treated with surgery alone or **(B)** surgery plus chemotherapy, according to CD44v6 status; **(C)** Kaplan-Meier estimates showing RFS of CD44v6_0 **(D)** CD44v6_1+, **(E)** CD44v6_2+ and **(F)** CD44v6_3+ GC patients, according to whether they were treated with surgery alone or with surgery plus chemotherapy; **(G)** Forest plot showing whether RFS is favored when treating patients with surgery plus chemotherapy or surgery alone, according to CD44v6 classification. No statistical significance was obtained, **(H)** Mean time to relapse of patients according to the CD44v6 sub-categories and treatment type. This analysis was carried out by exploring the CD44v6 sub-categories in the sub-set of patients that relapsed (n=73: 29 treated vs 40 untreated patients). The number of patients in each group is shown above the bars.

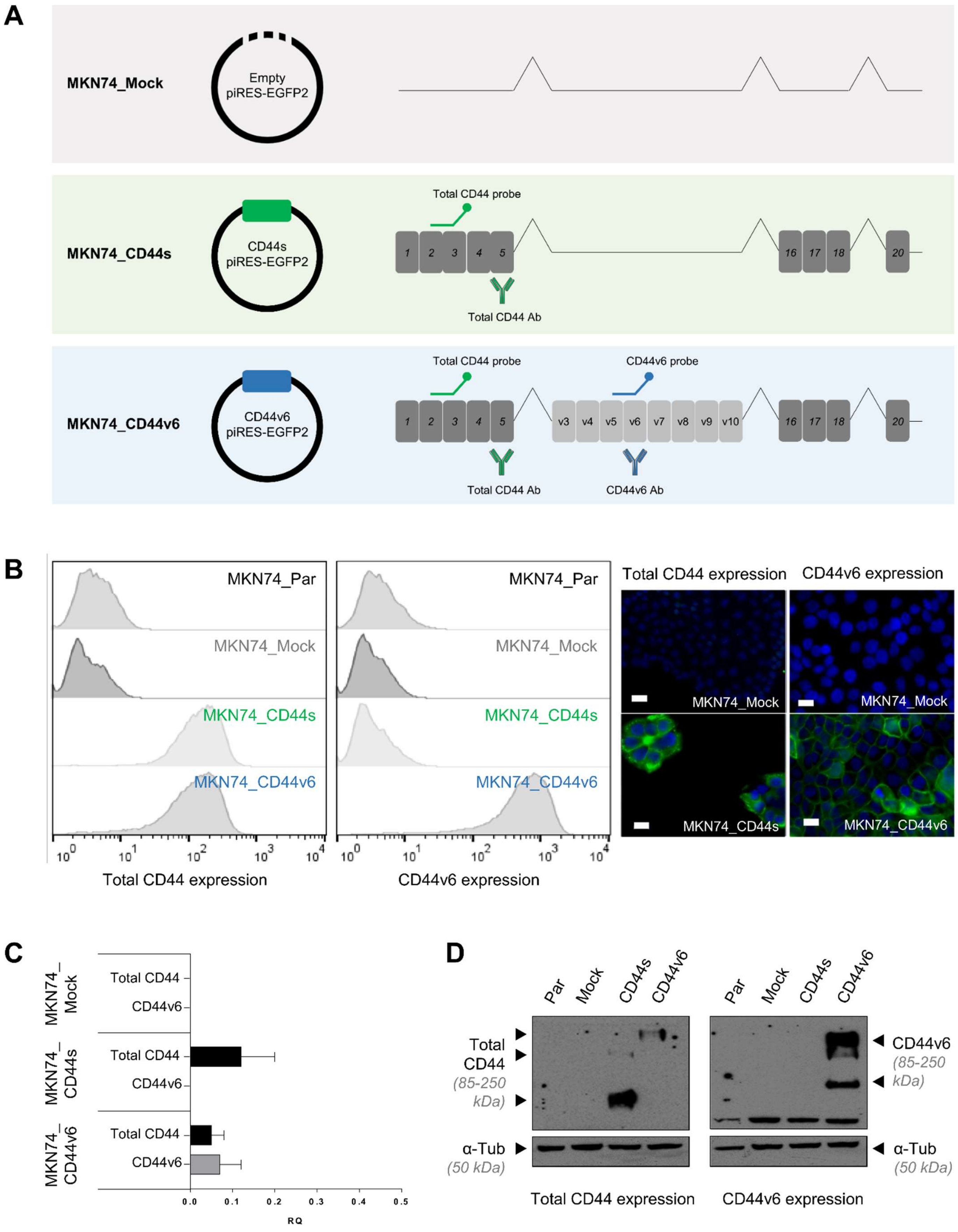
A
MKN74_Mock
Empty piRES-EGFP2
MKN74_CD44s
CD44s piRES-EGFP2
Total CD44 probe
Total CD44 Ab
MKN74_CD44v6
CD44v6 piRES-EGFP2
CD44v6 probe
CD44v6 Ab
B
MKN74_Par
MKN74_Mock
MKN74_CD44s
MKN74_CD44v6
Total CD44 expression
CD44v6 expression
C
Total CD44
CD44v6
RQ
D
Par
Mock
CD44s
CD44v6
Total CD44
(85-250 kDa)
α-Tub
(50 kDa)
Total CD44 expression
CD44v6 expression

**Appendix Fig 5:** The generated MKN74 cells express the intended CD44 transcripts. **(A)** Transcripts transfected into the MKN74 (CD44v6 negative) GC cell line. The CD44 canonical form is the long cytoplasmic tail, exon 20 containing isoform (CD44-03 – ENST00000263398). The exon v6 containing transcript expresses the variable exons between v3-v10 (variant CD44-04 – ENST00000415148). Ensembl release 68, July 2012. It is highlighted where the probes and antibodies, used below, bind to the cDNA or protein, respectively; **(B)** Expression was assessed at the post-translational level by immunofluorescence and flow cytometry against total CD44 and CD44v6. Nuclei are stained with DAPI (represented in blue) and white scale bars represent a distance of 20 μm; **(C)** Real time qRT-PCR representing the fold change expression of total CD44 and CD44v6 in transfected cells. Results are shown as average + SD and are representative of three independent experiments. The parental and MKN74_Mock transfected cells were used as negative controls; **(D)** Western Blot depicting total CD44 and CD44v6 protein levels in the generated MKN74 cells.

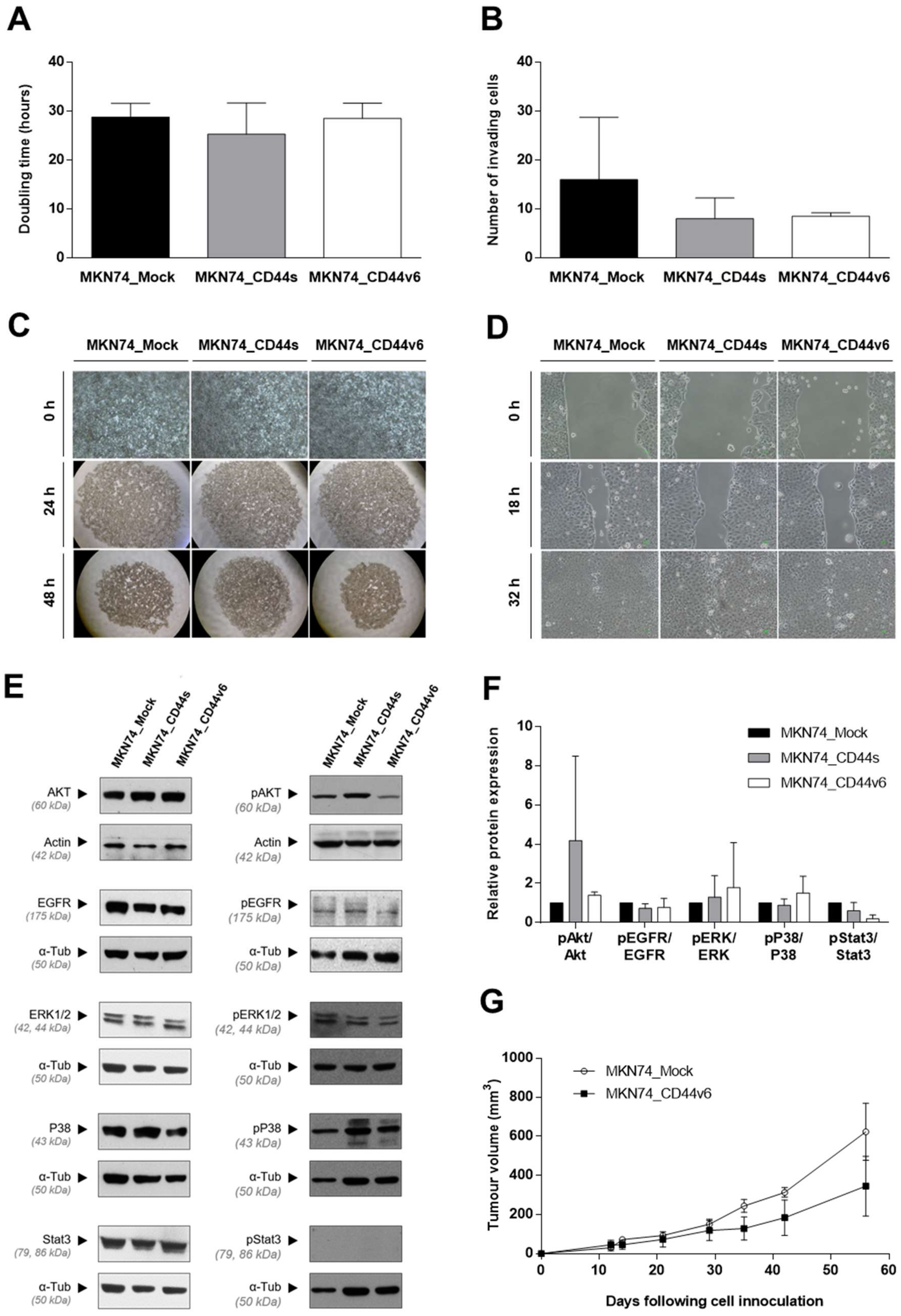

**Appendix Fig 6:** Functional characterization of the generated MKN74_Mock, MKN74_s and MKN74_CD44v6 cells regarding: **(A)** Doubling time; **(B)** Invasion capacity; **(C)** Slow aggregation assay; **(D)** Migration capacity; **(E)** Protein expression of common CD44 interactors: AKT, EGFR, ERK 1/2, P38 and Stat3 and respective phosphorylated forms; **(F)** Relative protein quantification of pAKT, pEGFR, pERK 1/2, pP38 and pStat3 with respect to the respective total protein amount. All data is represented as average + SD and/or are representative of three independent experiments; **(G)** Tumor growth kinetics of MKN74_Mock and MKN74_CD44v6 xenografts in nude mice, presented as average ± SEM. No statistical differences were observed.

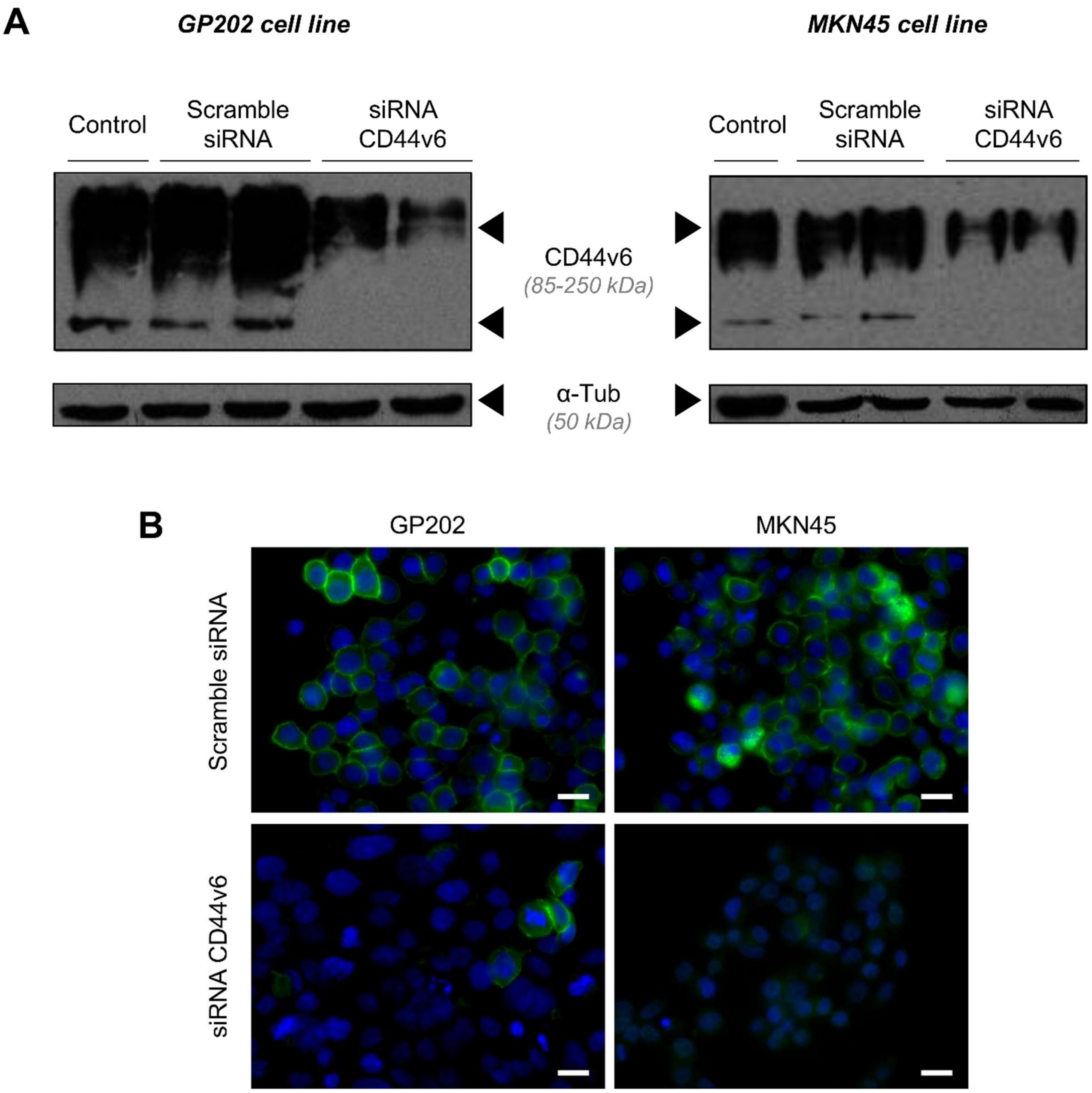

**Appendix Fig 7:** Confirmation of CD44v6 inhibition by siRNA in GP202 and MKN45 GC cell lines. **(A)** Western blotting of CD44v6 in GP202 and MKN45 upon incubation with scramble siRNA and CD44v6 siRNA; **(B)** Immunofluorescence of CD44v6 (represented in green) upon transfection with scramble siRNA and siRNA against CD44v6. Nuclei are stained with DAPI (represented in blue) and white scale bars represent a distance of 50 μm. Results are representative of three independent experiments.

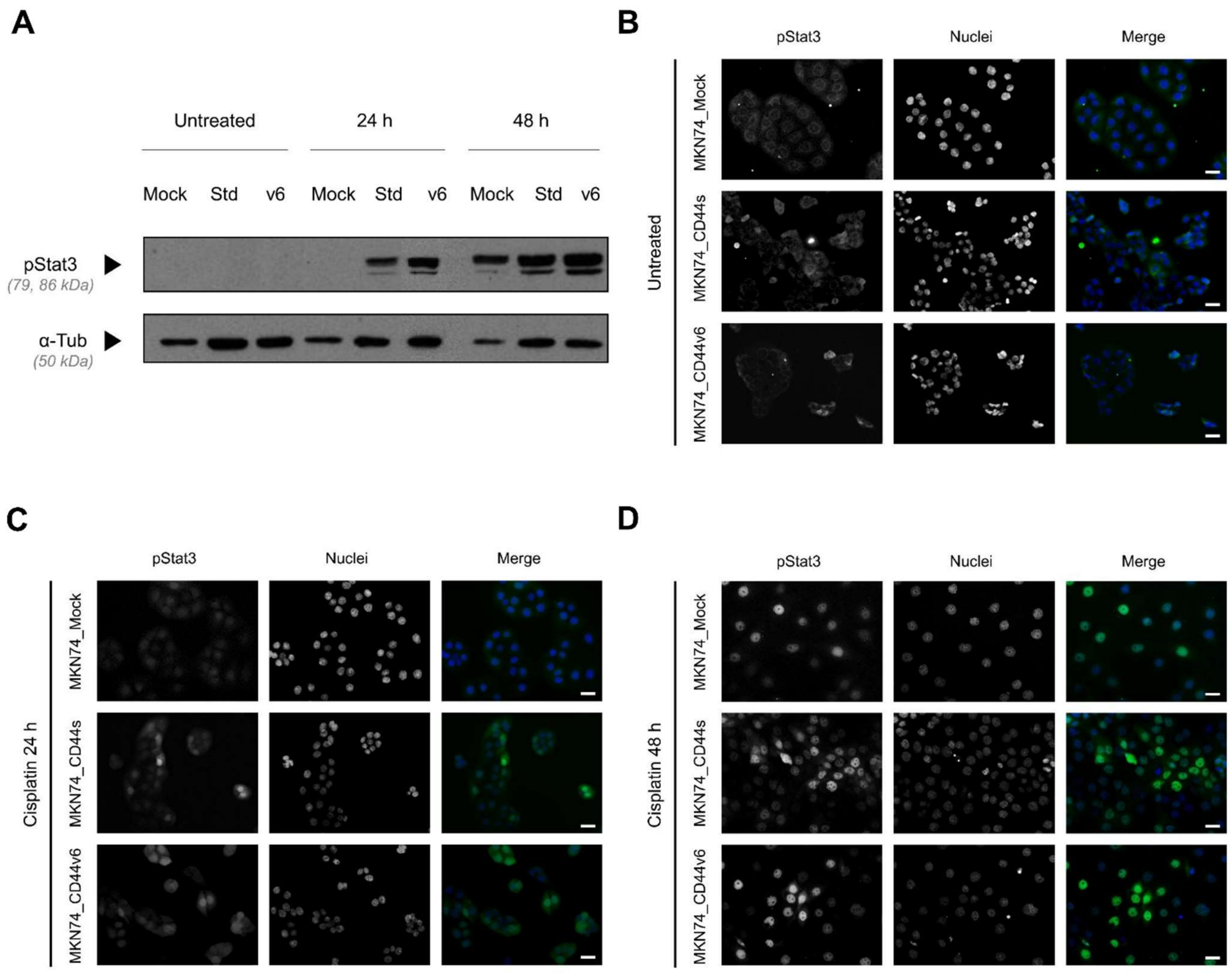

**Appendix Fig 8:** Early activation of Stat3 in CD44+ cells. **(A)** Western blotting of pStat3 in MKN-74 transfected cell lines; **(B)** Immunofluorescence of pStat3 (seen in green) in untreated cell lines (top right panel) and upon treatment with cisplatin for; **(C)** 24 hours (bottom left panel) and; **(D)** 48 hours (bottom right panel). Nuclei are stained with DAPI (seen in blue) and white scale bars represent a distance of 50 μm.

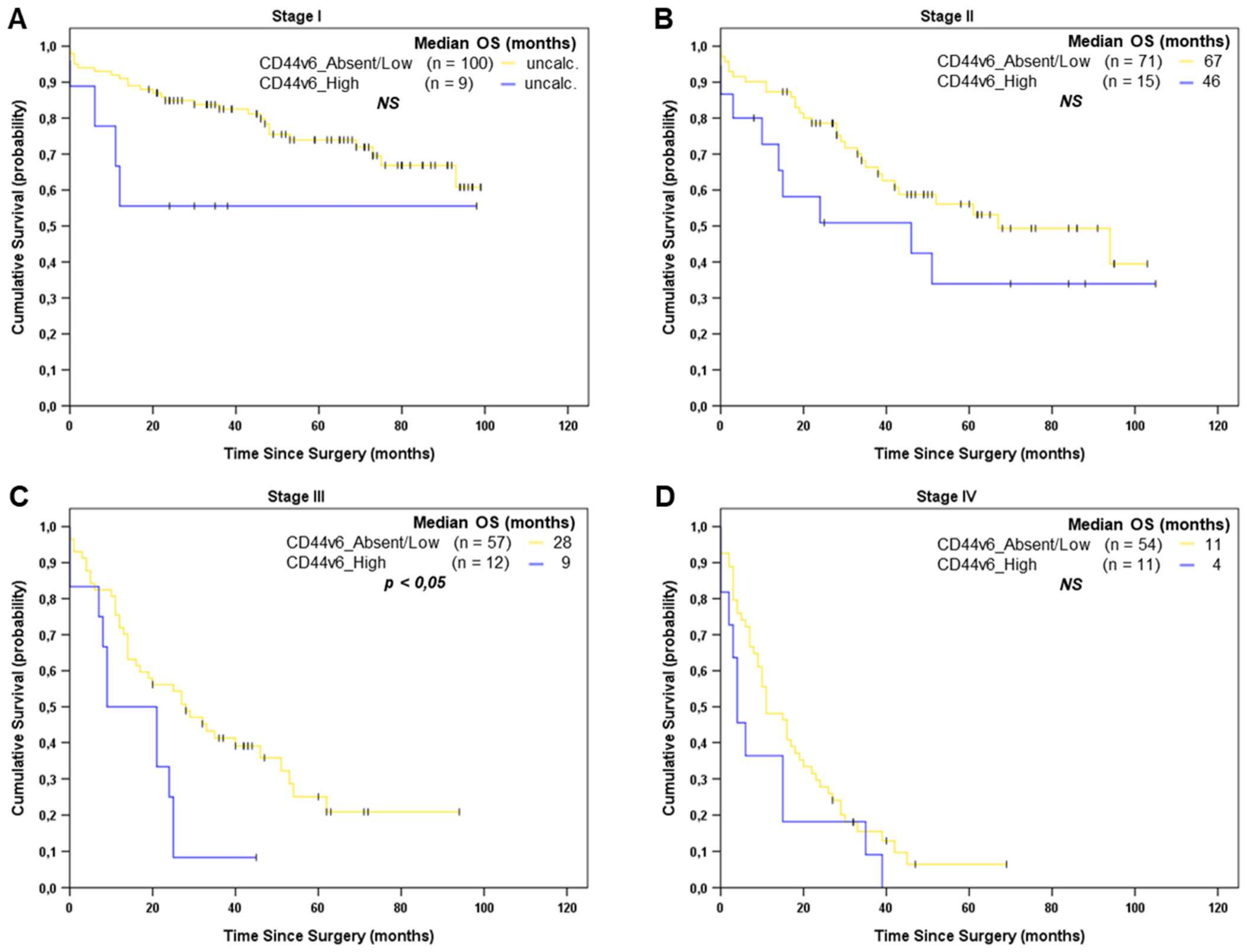

**Appendix Fig 9: (A)** Kaplan-Meier estimates showing OS in GC patients from TNM Stage I, **(B)** Stage II, **(C)** Stage III and **(D)** Stage IV, according to the CD44v6 status (CD44v6_absent/low vs CD44v6_high).

**Appendix Table 1:** Clinical-pathological characterization of the Gastric Cancer series.

| Variables | Total No. of patients (n=334) | | Surgery (n=203) | | Surgery + Chemotherapy (n=123) | |
|---|---|---|---|---|---|---|
| | No. | % | No. | % | No. | % |
| **Age, years** | | | | | | |
| Mean | 67.6 | | 71.02 | | 61.51 | |
| *Standard Deviation* | *11.9* | | *11.52* | | *9.89* | |
| **Gender** | | | | | | |
| Male | 188 | *56.3%* | 101 | *49.8%* | 82 | *66.6%* |
| Female | 146 | *43.7%* | 102 | *50.2%* | 41 | *33.3%* |
| Male to Female ratio | 1.3:1 | | 1:1 | | 2:1 | |
| **Lauren classification** | | | | | | |
| Intestinal | 156 | *46.7%* | 102 | *50.2%* | 50 | *40.7%* |
| Diffuse | 44 | *13.2%* | 20 | *9.9%* | 23 | *18.7%* |
| Mixed | 86 | *25.7%* | 51 | *25.1%* | 33 | *26.8%* |
| Indeterminate | 48 | *14.4%* | 30 | *14.8%* | 17 | *13.8%* |
| **WHO classification** | | | | | | |
| Tubular | 146 | *43.7%* | 97 | *47.8%* | 46 | *37.4%* |
| Papillary | 2 | *0.6%* | 2 | *1.0%* | 0 | *0.0%* |
| Mucinous | 7 | *2.1%* | 4 | *2.0%* | 3 | *2.4%* |
| Poorly cohesive | 38 | *11.4%* | 19 | *9.4%* | 18 | *14.6%* |
| Other types | 141 | *42.2%* | 81 | *39.9%* | 56 | *45.5%* |
| **Growth pattern** | | | | | | |
| Expansive | 61 | *18.3%* | 50 | *24.6%* | 10 | *8.1%* |
| Infiltrative | 259 | *77.5%* | 142 | *70.0%* | 110 | *89.4%* |
| Unclassified | 14 | *4.2%* | 11 | *5.4%* | 3 | *2.4%* |
| **Wall invasion** | | | | | | |
| Mucosa + Submucosa | 79 | *23.7%* | 70 | *34.5%* | 9 | *7.3%* |
| Muscular | 42 | *12.6%* | 29 | *14.3%* | 13 | *10.6%* |
| Subserosa + Serosa | 201 | *60.2%* | 100 | *49.3%* | 95 | *77.2%* |
| Other organs | 12 | *3.6%* | 4 | *2.0%* | 6 | *4.9%* |
| **Lymphatic permeation** | *332*[*1]* | | | | | |
| Absent | 103 | *31.0%* | 77 | *38.3%* | 25 | *20.3%* |
| Present | 229 | *69.0%* | 124 | *61.7%* | 98 | *79.7%* |
| **Perineural invasion** | *333*[*1]* | | | | | |
| Absent | 173 | *52.0%* | 125 | *61.6%* | 45 | *36.9%* |
| Present | 160 | *48.0%* | 78 | *38.4%* | 77 | *63.1%* |
| **Vascular invasion** | *331*[*1]* | | | | | |
| Absent | 138 | *41.7%* | 98 | *49.0%* | 40 | *32.5%* |
| Present | 193 | *58.3%* | 102 | *51.0%* | 83 | *67.5%* |
| **Surgical margins** | *333*[*1]* | | | | | |
| R0 | 298 | *89.5%* | 187 | *92.1%* | 104 | *85.2%* |
| R1/R2 | 35 | *10.5%* | 16 | *7.9%* | 18 | *14.8%* |
| **Depth of invasion (T)** | | | | | | |
| pT1 | 79 | *23.7%* | 70 | *34.5%* | 9 | *7.3%* |
| pT2 | 79 | *23.7%* | 49 | *24.1%* | 30 | *24.4%* |
| pT3-T4 | 176 | *52.7%* | 84 | *41.4%* | 84 | *68.3%* |
| **Lymph node metastases (N)** | *333*[*1]* | | | | | |
| Absent (pN0) | 129 | *38.7%* | 113 | *55.9%* | 15 | *12.2%* |
| Present (pN+) | 204 | *61.3%* | 89 | *44.1%* | 108 | *87.8%* |
| **Distant metastases (M)** | | | | | | |
| Absent | 265 | *79.3%* | 176 | *86.7%* | 88 | *71.5%* |
| Present | 69 | *20.7%* | 27 | *13.3%* | 35 | *28.5%* |
| **pTNM Staging** | | | | | | |
| I | 109 | *32.6%* | 101 | *49.8%* | 8 | *6.5%* |
| II | 87 | *26.0%* | 50 | *24.6%* | 36 | *29.3%* |
| III | 69 | *20.7%* | 25 | *12.3%* | 44 | *35.8%* |
| IV | 69 | *20.7%* | 27 | *13.3%* | 35 | *28.5%* |
| **Disease recurrence** | 265[*2] | | | | | |
| Absent | 192 | *72.5%* | 144 | *81.8%* | 47 | *53.4%* |
| Present | 73 | *27.5%* | 32 | *18.2%* | 41 | *46.6%* |

[*1] Remaining data not available; [*2] Stage IV patients were excluded; [*3] These pTNM stage I patients received chemotherapy because they had been considered as locally advanced at diagnosis; [*4] These pTNM stage IV patients did not receive chemotherapy due to comorbidities/general fitness. pTNM (pathological tumor-node-metastasis).

**Appendix Table 2:** Platinum (Pt) and non-Pt based chemotherapeutic regimens administered to patients in this GC cohort.

| | Chemotherapeutic Regimen | No. of Patients per regimen |
|---|---|---|
| Platinum-based | Cisplatin + 5-FU + Taxotere | 27 |
| | Cisplatin + Capecitabine | 26 |
| | Cisplatin + 5-FU | 9 |
| | Cisplatin + 5-FU + Docetaxel | 6 |
| | Cisplatin + 5-FU + Etoposide | 2 |
| | Cisplatin + Capecitabine + Taxotere | 2 |
| | Cisplatin + 5-FU + Epirubicin | 2 |
| | Carboplatin + 5-FU + Taxotere | 3 |
| | Carboplatin + Docetaxel | 1 |
| | Oxaliplatin + Capecitabine | 17 |
| | Oxaliplatin + Capecitabine + Epirubicin | 17 |
| | Oxaliplatin + 5-FU + Leucovorin | 1 |
| | Capecitabine | 6 |
| | Leucoviron + 5-FU + irinotecan | 2 |
| | Leucovorin + 5-FU | 1 |
| | Irinotecan + Capecitabine | 1 |
